# Organizing principles in the nitrogen–carbon landscape of marine heterotrophic bacteria

**DOI:** 10.64898/2026.06.04.730220

**Authors:** Franziska Kratzl, Helen Scott, Simran Jayasinghe, Kendall Hughes, Melisa Osborne, Daniel Sher, Daniel Segrè

**Affiliations:** Bioinformatics Program, Faculty of Computing and Data Science, Boston University, Boston, MA 02215, USA; Department of Biology, Boston University, Boston, MA 02215, USA; Biological Design Center, Boston University, Boston, MA 02215, USA; Program in Biochemistry, Smith College, Northampton, MA 01063; Department of Marine Biology, Leon H. Charney School of Marine Sciences, University of Haifa, Haifa 3498838, Israel; Department of Physics, Boston University, Boston, MA 02215, USA; Department of Biomedical Engineering, Boston University, Boston, MA 02215, USA

## Abstract

Marine microbes metabolize a wide range of carbon and nitrogen sources, shaping global biogeochemical cycles. Despite being crucial at the global scale, the coupling between carbon and nitrogen remains poorly understood at the level of individual metabolites and bacteria. By phenotyping a library of marine heterotrophic bacteria across increasingly complex carbon and nitrogen sources, we generated a snapshot of this coupling. Growth phenotypes were weakly explained by phylogeny, but could be organized around substrate properties, including C:N stoichiometry and degree of reduction. Beyond explainable average trends along these axes, different strains displayed significant variability. For some bacteria, nitrogen use efficiency was associated with ammonium secretion, consistent with their role in ecological interactions. Furthermore, nonlinearities emerged in how yields depend on combinations of resources, such as mixtures of amino acids and dipeptides. These organizing principles may help understand the ecological role of marine heterotrophic bacteria and parametrize models of biogeochemical cycles.

## Introduction

Microorganisms drive the global nitrogen cycle (N-cycle) through the assimilation, transformation, and remineralization of nitrogen-containing compounds [1–4]. In the oceans, this processes connect the dissolved organic nitrogen (DON) pool with the dissolved inorganic nitrogen (DIN) pool, converting complex organic molecules into inorganic forms and back again. Heterotrophic bacteria, increasingly recognized as key players in marine biogeochemical processes, are major drivers of these transformations [5–7]. While DIN species are well defined (NH_4_^+^, NO_3_, NO_2_ ^–^), the DON pool is chemically diverse and still poorly characterized, ranging from low-molecular-weight compounds such as amino acids, vitamins, and other specialized metabolites to high-molecular-weight molecules such as peptides [8–10]. Available nitrogen is frequently assimilated directly into biomass, as extensive dedicated nitrogen storage pools beyond biomass-associated nitrogen are uncommon in many heterotrophic bacteria [11, 12]. Consequently, in marine environments where nitrogen is often limiting, the choice of nitrogen source can strongly influence nutrient recycling and microbial interactions such as competition and cross-feeding [13–16]. This complexity is further increased by the fact that DON molecules can serve as nitrogen sources, carbon sources, or both, requiring diverse metabolic pathways. The assimilated carbon and nitrogen can then be allocated either to biomass production or to energy generation.

Despite the importance of disentangling this complexity, heterotrophic bacterial phenotypes are often studied primarily through the lens of carbon utilization [17, 18], and covalently bound C–N molecules are seldom analyzed as both carbon and nitrogen sources. However, microorganisms encounter environments with diverse elemental stoichiometries that determine the limiting nutrient and thereby influence growth strategies [19]. For example, environmental C:N ratios shift towards lower values in protein-rich environments, as proteins are relatively nitrogen-rich compounds [13, 20, 21]. While co-consumption and prioritization of co-occurring substrates in heterotrophic bacteria have been studied in specific cases [22, 23], the broader question of how different heterotrophic bacteria process diverse DIN (in combination with carbon sources) and DON molecules remains open.

Here, we address this question by generating and analyzing a large dataset to systematically capture how diverse marine heterotrophs utilize different combinations of carbon (C), nitrogen (N), and combined C–N sources of increasing chemical complexity (see Figure 1**A**). We screened growth across more than 50 defined media, generating over 6,000 growth curves with a focus on both carbon and nitrogen supply. These experiments were performed for a marine bacterial library comprising 42 heterotrophic strains representing major lineages from oligotrophic and temperate ocean environments [17, 24]. We thus derived a phenotypic landscape of carbon and nitrogen utilization.

**Figure 1:**
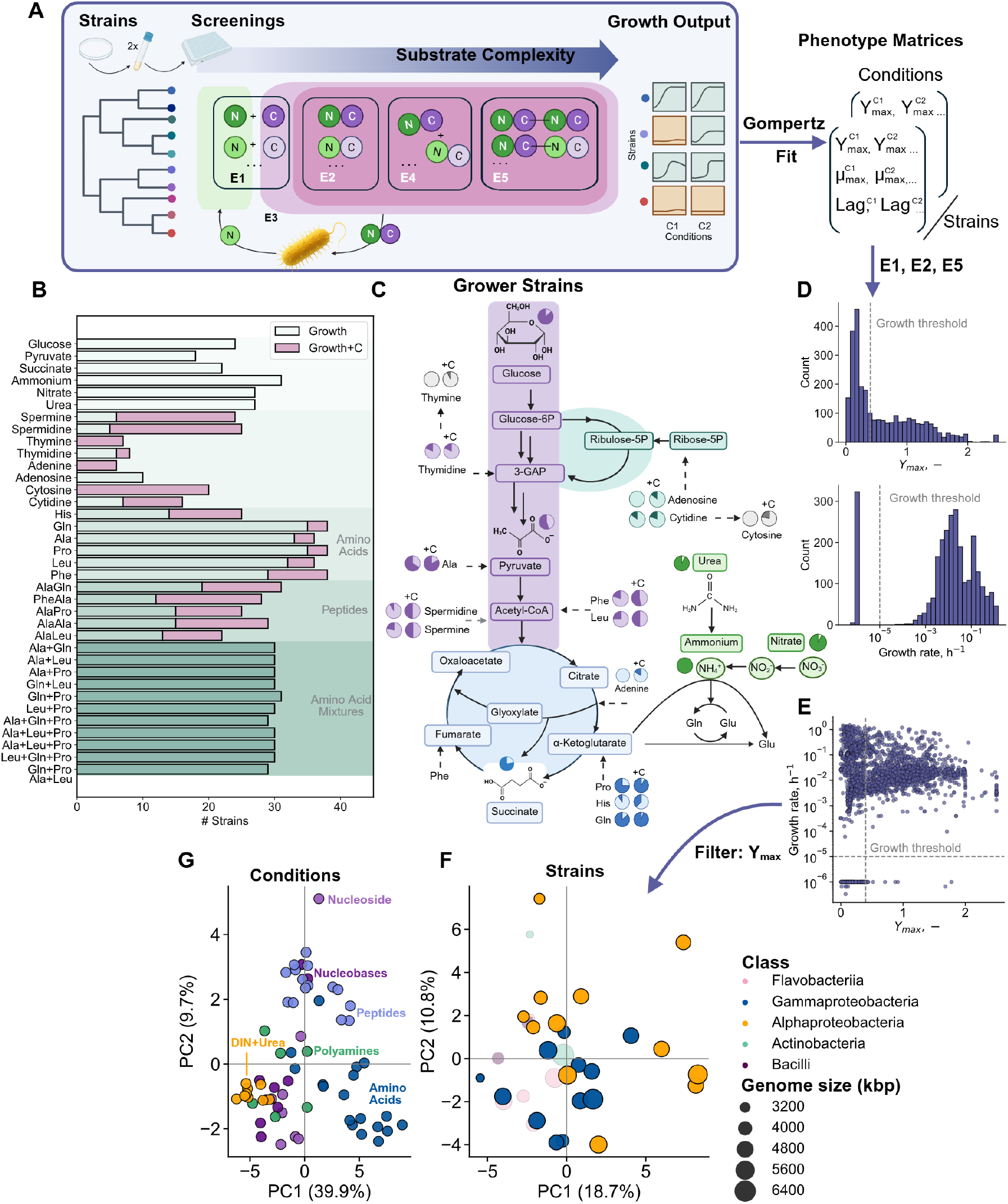
Experimental overview and quantitative phenotyping framework for mapping microbial growth across nitrogen–carbon landscapes. **(A)** Overview of experiments E1–E5. A total of 42 marine heterotrophic strains were screened for growth on combinations of simple carbon and nitrogen substrates (E1) and increasingly complex dissolved organic nitrogen (DON) compounds, including amino acids, polyamines, nucleobases, nucleosides, amino acid mixtures, and dipeptides (E2–E5). Circle colors indicate carbon- and nitrogen-containing molecular moieties. **(B)** Number of strains growing across substrates. **(C)** Number of strains classified as growers, shown together with the predicted entry points of substrates into central metabolism. +C indicates growth on molecule plus additional carbon source. **(D)** Growth curves were fitted using a Gompertz function to generate phenotype matrices containing maximal biomass accumulation (*Y*_max_), maximal growth rate (*µ*_max_), and lag phase (*Lag*) for each strain and condition. Histograms show the distributions of growth rate and yield across experiments E1, E2, and E5. **(E)** Weak nonlinear relationship between maximal growth rate and maximal biomass accumulation across conditions from E1, E2, and E5. **(F)** Principal component analysis (PCA) of yield phenotypes shows only weak separation between phylogenitc classes (Eg. Gammaproteobacteria and Alphaproteobacteria along PC1 and PC2.) **(G)** PCA of the transposed phenotype matrix reveals clustering by substrate condition.

Beyond characterizing nutrient utilization, we searched for general patterns that could link microbial metabolism to biogeochemical processes. We first show that the phenotype yields are better explained by substrate classes then by phylogeny. Motivated by this observation, we next asked whether simple properties of substrates could explain the observed phenotypic variation. Specifically, we examined how substrate C:N ratio, which determines the balance between carbon and nitrogen supply, and degree of reduction, which reflects the energetic content of a substrate, shape microbial growth phenotypes. Using both experimental measurements and genome-scale metabolic models, we found that these properties can organize the biomass yields and growth, providing a mechanistic link between substrate chemistry and microbial metabolism. Combined with measurements of ammonium release during growth on amino acids, this framework allowed us to estimate nitrogen use efficiency (NUE) and relate metabolic behavior to the ecological roles of two strains in greater detail. Finally, we explored how nutrient context, eg. through mixtures of amino acids, influences microbial phenotypes and can generate strain-specific nonlinear growth responses. Overall, beyond providing a large phenotypic dataset, our work highlights key aspects of carbon–nitrogen coupling in marine heterotroph metabolism and proposes organizing principles that may help improve ecological models of microbial communities.

## Results

### Generating a phenotypic landscape of nitrogen and carbon utilization across heterotrophic bacteria

To generate a dataset on nitrogen, carbon and nitrogen-carbon use across 42 marine heterotrophic bacteria, we performed multiple metabolic screenings (see Figure 1**A** E1-E5). All screenings were initiated from single colonies, transferred to marine broth, and subsequently acclimated in nitrogen-starvation medium prior to growth in 96-well plates for 1–2 weeks (see Methods for details).

In experiment E1 (see Figure 1**A**), we tested three separate nitrogen sources (nitrate, ammonium, urea) and three central carbon sources (glucose, pyruvate, succinate), resulting in nine carbon and nitrogen combinations (DIN+Urea). Examining growth pattern across these conditions revealed that approximately 75% of strains grew in at least one condition (E1-growers), whereas the remaining strains showed no growth across all tested conditions (Figure 1**B & C**). The amount of non-growers is consistent with previous similar metabolic profiling studies (see Supplement Table 2) [17]. Among E1-growers, all strains utilized ammonium, while four strains failed to grow on nitrate or urea (see Figure 1**C** and Figure S1). Consistent with these phenotypes, roughly 25% of genomes lacked annotated nitrate or nitrite reduction genes, and many of these additionally lacked urease genes, suggesting specialization toward reduced or more complex organic nitrogen sources (see Supplement Table 3 and Figure S2). This pattern was pronounced in Flavobacteria which are known to degrade complex biopolymers [25]. The usability of carbon sources was more variable (see Figure S1): 13 strains failed to grow on pyruvate, seven on glucose, and nine on succinate. Notably, pyruvate utilization could not be reliably predicted from genomic annotations, as transporter genes were inconsistently annotated across genomes (see Supplement Table 4, 5 and Figure S3). Some strains showed reproducible growth without an added nitrogen source, most notably *Rhodococcus erythropolis* 100887, consistent with previous reports of a closely related extreme oligotroph capable of utilizing trace amounts of atmospheric ammonia [17, 26]. In total, we identified 11 strains with similar putative extreme “nitrogen oligotrophic” behavior, prompting us to spatially separate negative controls from experimental conditions in all subsequent assays to minimize air mediated ammonia cross-feeding between adjacent wells (see Supplement Table 6).

In Experiment E2, we expanded the screening to chemically diverse dissolved organic nitrogen (DON) compounds, including amino acids, polyamines, nucleosides, and nucleobases spanning a broad substrate C:N stoichiometry, ranging from 1 to 9 (Figure 1**B & C**). Because carbon and nitrogen are intrinsically coupled within DON molecules, substrates were supplied either alone or supplemented with additional carbon (pyruvate and glucose) or nitrogen (ammonium) sources to probe nutrient (elemental) limitation effects. Alanine, glutamine, leucine, and proline supported growth in the largest number of strains as sole nitrogen and carbon sources (Figure 1**C**). While growth on histidine alone was rare, many strains utilized it as a nitrogen source when supplemented with additional carbon (Supplement Figure 1**B**). Notably, all non-growers from E1 grew on at least one amino acid (Supplement Table 2). In contrast, growth on polyamines as sole carbon and nitrogen sources was uncommon, although many strains utilized them when supplemented with additional carbon. Similarly, none of the tested nucleobases supported growth alone, whereas several strains grew on the corresponding nucleosides (Figure 1**B**; Supplement Figure S4 & S5), consistent with recent work suggesting that heterocyclic nitrogen compounds may represent relatively inert or “dormant” components of the DON pool [27].

To investigate how increasing molecular complexity influences growth, we first extended our screening to amino acid mixtures, encompassing all possible pairs, triplets and quadruplets of alanine, glutamine, proline, and leucine, while maintaining a constant total nitrogen supply (see Figure 1**A**, Experiment E4). These mixtures supported growth in a large and similar fraction of strains, potentially because distinct amino acids within the mixtures could differentially satisfy carbon and nitrogen demands (see Figure 1**B** and Supplement Figure S6). We increased substrate complexity further by screening five alanine-containing dipeptides as combined carbon and nitrogen sources (Experiment E5). Individual dipeptides supported growth in 15–20 strains, and additional carbon supplementation further increased the number of strains able to grow (see Supplement Figure S7).

Overall, these screenings led to the generation of multiple quantitative phenotype matrices describing growth of all strains across chemically diverse nutrient environments, amounting to what we refer to as a carbon-nitrogen phenotype landscape Each matrix contains three growth parameters associated with each experiment: lag phase (*Lag*), maximal growth rate (*µ*_*max*_), and maximal optical density (*Y*_*max*_). These parameters were estimated by fitting a Gompertz function to the growth curves (Figure 1**A&E**; Supplement Figure S8) [28]. The distributions of *Y*_*max*_ and *µ*_*max*_ across experiments E1, E2, and E5 are shown in Figure 1**D**. Growth rate and maximal biomass accumulation showed only a weak non-linear association across conditions (Figure 1**E**). Equipped with these data we started analyzing the phenotypes in search for organizing principles that would help us better understand the observations and their ecological implications.

### Microbial growth phenotypes show weak association with phylogeny

Since closely related strains may share similar nutrient preferences, we first asked whether growth phenotypes could be explained by phylogeny or by the supplied carbon and nitrogen sources. In particular, we focused on analyzing the inferred yield (*Y*_*max*_) across screenings E1, E2, and E5 (see Supplement Figure S9). Principal component analysis (PCA) revealed only weak clustering by phylogeny, indicating that overall growth phenotypes were largely decoupled from taxonomic relationships (Supplement Figure 1**F**). Nevertheless, among the five bacterial phylogenetic classes represented in the screenings, the mean of Alphaproteobacteria and Gammaproteobacteria showed significant separation along PC2 (p=0.012; not significant on PC1). The primary drivers underlying this separation were growth on peptides and amino acids, suggesting that Alphaproteobacteria reached, on average, slightly higher optical densities on these C–N sources (see Supplement Figure S9). It has been shown that peptide degradation can vary considerably among bacteria within the same taxonomic class relating to nutrient partitioning [29]. Neither genome size nor GC content explained the observed phenotype structure. Likewise, inclusion of additional ecological metadata such as isolation depth and sampling location did not reveal stronger organization within the dataset across phylogenetic traits (Supplement Figure S9). The limited phylogenetic structure observed here may reflect the fact that many substrates we tested belong to the conserved core metabolism of heterotrophic bacteria, as shown in similar studies [17, 18].

In contrast to the PCA by strains, substrate classes (conditions) showed substantially stronger clustering (Figure 1**G**). For example, phenotypes associated with amino acids form distinct clusters from those observed for DIN+urea conditions, while peptide-associated phenotypes cluster closer to the amino acid phenotypes. This pattern suggests that substrate exerts a stronger influence on growth phenotypes than phylogeny. Motivated by this result, we next focused on analyzing our data in more depth from the perspective of substrates, with special focus on key molecular properties linked to the stoichiometric or energetic constraints that shape growth [19, 30]. In particular, we revisited the growth phenotypes as a function of carbon-to-nitrogen (C:N)-ratio and redox-state, as well as substrate context (e.g., mixtures of molecules), to identify organizing principles which may link the phenotypes to nutrient-use efficiency, substrate interactions, and ecological strategies across marine heterotrophs (Figure 2**A**).

**Figure 2:**
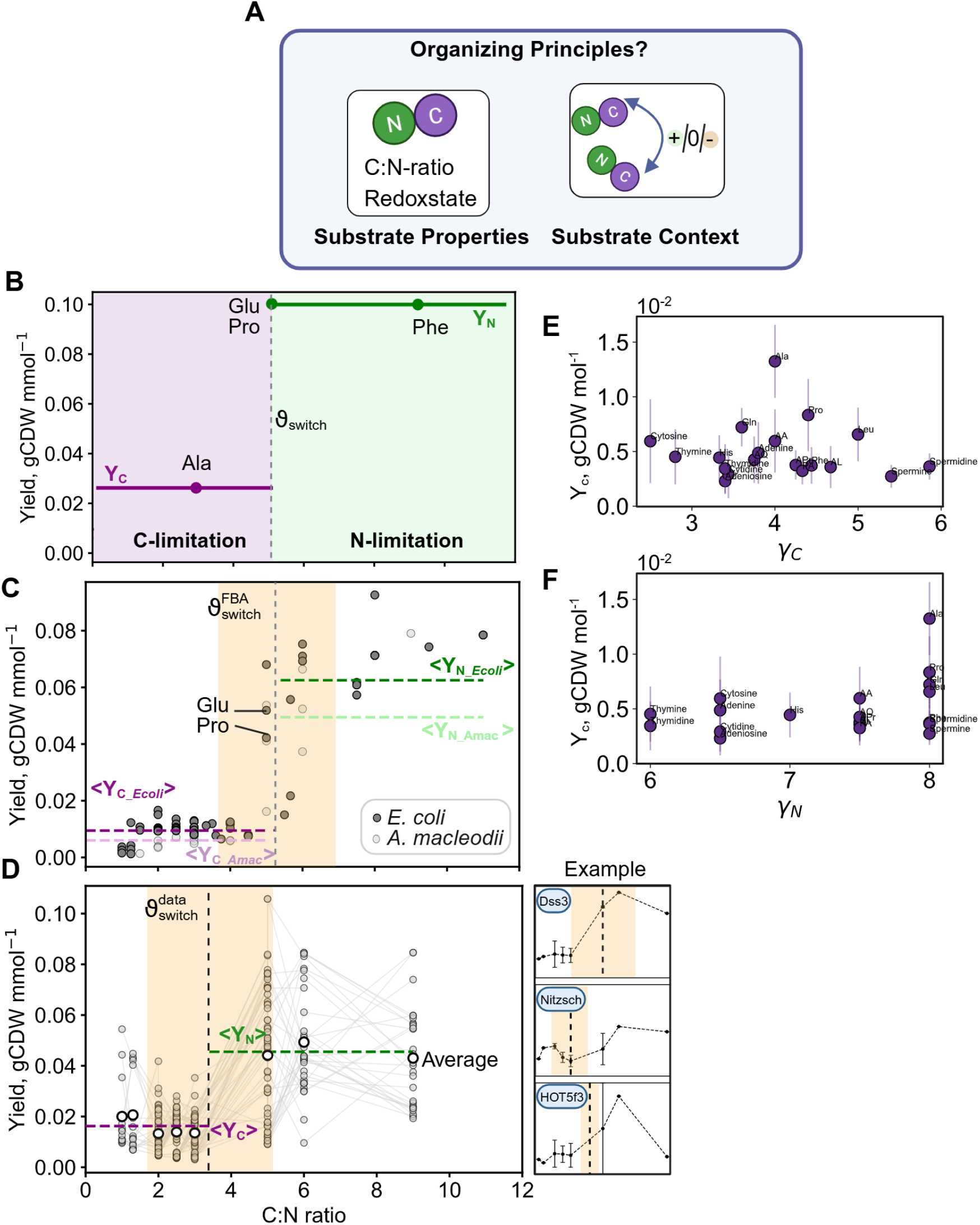
Substrate stoichiometry and degree of reduction organize microbial biomass yields. **(A)** Substrate properties and substrate context as potential organizing principles linking nutrient accessibility, substrate-use efficiency, and metabolic interactions across the phenotype landscape.**(B)** Conceptual yields for DON substrates across varying C:N ratios, showing a transition from carbon to nitrogen limitation at *ϑ*_switch_. Example yields are shown for four amino acids spanning a range of C:N ratios. *Y*_*C*_ = yield per limiting carbon; *Y*_*N*_ = yield per limiting nitrogen. **(C)** Simulations using genome-scale metabolic models (*E. coli* iML1515 and *A. macleodii* iHS4156) grown on DON substrates (upper bound: 10 mmol N gCDW^−1^ h^−1^). Curves show mean relationships between substrate C:N ratio and normalized yields. Glutamate and proline are highlighted (Glu = glutamate). Dashed lines are mean < *Y*_*C*_ *>* and < *Y*_*N*_ *>*. **(D)** Experimentally determined yields normalized by carbon (*Y*_*C*_) or nitrogen (*Y*_*N*_) across all strains and three representative examples. Conditions without growth were excluded. Smoothed curves and dashed lines indicate average yields under carbon-imited (C:N < 3.4) and nitrogen-limited (C:N ≥ 3.4) regimes. White circles indicate mean values at each C:N ratio. Degree of reduction as a substrate property based on [30]. Carbon-normalized yields (*Y*_*C*_) plotted against substrate-carbon degree of reduction (*γ*_*C*_) **(E)** and substrate-nitrogen degree of reduction (*γ*_*N*_). Points represent mean yields across all strains; error bars indicate standard deviations.

### Substrate C:N-ratio and degree of reduction constrain phenotypes

The elemental stoichiometry in a substrate has a central role because it determines how much carbon or nitrogen is available. A key aspect of providing only DON molecules is that carbon and nitrogen are supplied in a fixed proportion within each molecule, allowing us to systematically explore how bacteria respond to substrate C:N ratios that deviate from their biomass C:N demand. Using the endpoint of each growth curve (*Y*_*max*_), and assuming full substrate utilization, biomass yields can be expressed relative to substrate carbon (*Y*_*C*_) and nitrogen (*Y*_*N*_). When plotted against the substrate C:N ratio (denoted *ϑ*), two regimes are expected (Figure 2**B**): below a threshold *ϑ*_switch_, carbon is limiting and biomass increases proportionally with carbon, resulting in a constant *Y*_*C*_. Above this threshold, nitrogen becomes limiting and additional carbon no longer increases biomass, resulting in a constant *Y*_*N*_ . *ϑ*_switch_ is expected to be higher than the biomass C:N ratio (*ϑ*_Biomass_), as some carbon is lost to respiration and maintenance (Supplementary Text S11). In principle, the value of *ϑ*_switch_ can be inferred from the data based on the theoretical result that *ϑ*_switch_ = *Y*_*N*_ */Y*_*C*_ (see Supplementary Methods). In practice, however, metabolic noise limits the reliability of this approach. In the analysis below, we therefore use an alternative method to estimate *ϑ*_switch_ for simulated and experimental data (see Supplement Text S11 and Figure S12).

Figure 2**C** shows yield predictions from the *E. coli* genome-scale metabolic model across substrates with varying C:N ratios, providing a theoretical benchmark for the observed relationship (Supplementary Table 7). Genome-scale metabolic models enable calculations of cellular carbon and nitrogen budgets based on detailed reaction stoichiometry. Here, we used such models to capture the overall C and N balance and compute the yields (*Y*_*C*_, *Y*_*N*_). Specifically, we used the model of *A. macleodii* MIT1002 iHS4156 (v3.0.0, manuscript in preparation) and the well-curated *E. coli* model iML1515 to simulate growth on substrates provided as sole carbon and nitrogen sources using flux balance analysis (FBA; see Supplement). Simulated growth rates can be converted to yields by normalizing with respect to the uptake flux of the limiting element. FBA calculations for *E. coli* predict a 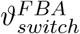 of 5.2 ± 1.3 (see Figure 2**D**, Supplement Text 11), and display the expected constant yields relative to C at low C:N ratios (*Y*_*C*_ = 0.01 ± 0.003 gCDW mmol^−1^), and relative to N at high C:N ratios (*Y*_*N*_ = 0.063 ± 0.02 gCDW mmol^−1^). A noisier, but similar relationship can be seen for *A. macleodii* (*Y*_*C*_ = 0.006 ± 0.002 gCDW mmol^−1^, *Y*_*N*_ = 0.05 ± 0.021 gCDW mmol^−1^). Notably, while the simulation results follow the expected trend, they vary across both strains and substrates. Substrates with identical C:N ratios (e.g., proline and glutamate) can yield different outcomes, reflecting substrate-specific biochemical constraints captured by metabolic models, such as differences in ATP or NAD(P)H requirements.

Actual experimental growth on individual DON molecules as sole carbon and nitrogen sources (experiment E2) allows us to test the relevance of this C:N organization principle across diverse marine heterotrophs. It is not obvious that 42 strains grown on 13 DON molecules would exhibit behavior consistent with this principle. Individual bacteria may respond differently to specific metabolites, vary in their biomass C:N composition, and may not fully or uniformly utilize the available substrates under the tested conditions. Despite this variability, averaging across strains and substrates (Figure 2**D**) reveals a pattern consistent with that observed in the simplified cases (Figure 2**D**).

In particular, the averages of Y_*C*_ and Y_*N*_ across all strains can be used to estimate the C:N ratio at which the limitation switches from C to N (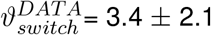, see Supplementary Text S11 and Figure S12). As shown in Figure 2**D**, for 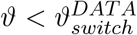, i.e. under carbon-limited conditions, the average Y_*C*_ depends very weakly on *ϑ* (slope = -0.004, p = 0.054) and has an average value Y_*C*_ = 0.014 ± 0.009 gCDW mmol^−1^, which is comparable to the yields predicted by FBA. For 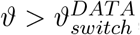, i.e. under nitrogen-limited conditions, the yield per nitrogen is approximately constant, with a value Y_*N*_ = 0.046 ± 0.020 gCDW mmol^−1^ (slope = -0.002, p = 0.7). This value agrees well with the predicted yields of the *Alteromonas* model but is substantially lower than those predicted by the *E. coli* model (see Supplement Figure S12 and Table 7). One surprising feature of this analysis is the low mean value of 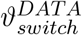 across strains (though the wide error bar makes any final interpretation very challenging). Given the fact that this value is expected to be an upper bound for the C:N ratio in the biomass, the implication would be a low average biomass C:N ratio across all heterotrophs (see Supplement Figure S12), in contrast with numbers typically reported in biotechnologically used strains such as *E. coli* (C:N 4-6) or ecological stoichiometry (marine bacterial C:N 4-6, soil bacteria 3-8) [31–33]. Interestingly, however, the curves (and corresponding *ϑ*_*switch*_) for individual strains display high variability. While for several individual strains, the yield graph is very noisy, some strains display a pattern that is highly consistent with the idealized expectation.

Our single-substrate design imposes defined stoichiometric constraints, enabling direct comparison between substrate-driven C:N variation and natural variability in marine environments, such as regional ocean gradients and localized microenvironments like marine snow [19, 34–36]. Notably, variability in yields differs between limitation regimes: carbon-limited yields are more constrained, whereas nitrogen-limited yields show greater variability across strains (see Supplementary Figure S12). This further suggests that strains differ substantially in how efficiently acquired nitrogen is retained versus released back into the environment.

The DON molecules impose a fixed carbon-to-nitrogen supply ratio that determines whether growth is carbon- or nitrogen-limited, giving rise to the observed transition at *ϑ*_switch_. However, substrate stoichiometry is not the only property expected to influence microbial growth as previously described [30]. DON substrates also differ in their energetic content, which can be quantified by their degree of reduction and reflects the average electron content per carbon (*γ*_*C*_) or nitrogen atom (*γ*_*N*_) (see Figure 2**E**). While *γ*_*C*_ is related to the energy potentially available during catabolism, *γ*_*N*_ reflects the energetic investment required for nitrogen assimilation into biomass (See Supplement S10). Across strains, carbon-normalized yields exhibited a non-monotonic relationship with *γ*_*C*_, with maximal yields observed at intermediate values, consistent both with classical thermodynamics derivations and with recent theoretical results for environmental microbes [37]. In contrast, substrates containing more oxidized nitrogen tended to support lower biomass yields (Figure 2**F**), consistent with the additional energetic costs associated with reducing nitrogen prior to incorporation into biomass. Note, that a mean degree of reduction calculated across carbon and nitrogen atoms showed a similar pattern to that observed for *γ*_*C*_ (Supplementary Figure S10).

### Nitrogen use efficiency in individual strains may shape micro-bial interactions

Under carbon limitation, when organic nitrogen is in excess, cells release surplus nitrogen via regulated processes or metabolic overflow, converting organic nitrogen into inorganic ammonium (mineralization) [38–40]. This process transfers nitrogen from the DON to the DIN pool and can shape nutrient availability within microbial communities. Nitrogen use efficiency (NUE) provides a quantitative framework to capture this behavior by describing the fraction of acquired nitrogen retained in biomass versus released (Figure 3**A**, see Supplement Text S13). Accordingly, high NUE reflects efficient nitrogen retention, whereas low NUE indicates nitrogen loss through mineralization. Variation in NUE may therefore describe differences in microbial interactions. For example, differential effects of bacteria in our collection (*Alteromonas* spp. and *Marinovum*) on *Prochlorococcus* in co-culture have been attributed to cross-feeding, including nitrogen exchange [41, 42]. Consistent with genome-scale model predictions, secretion of ammonium during growth on organic nitrogen may be metabolically favorable [43].

**Figure 3:**
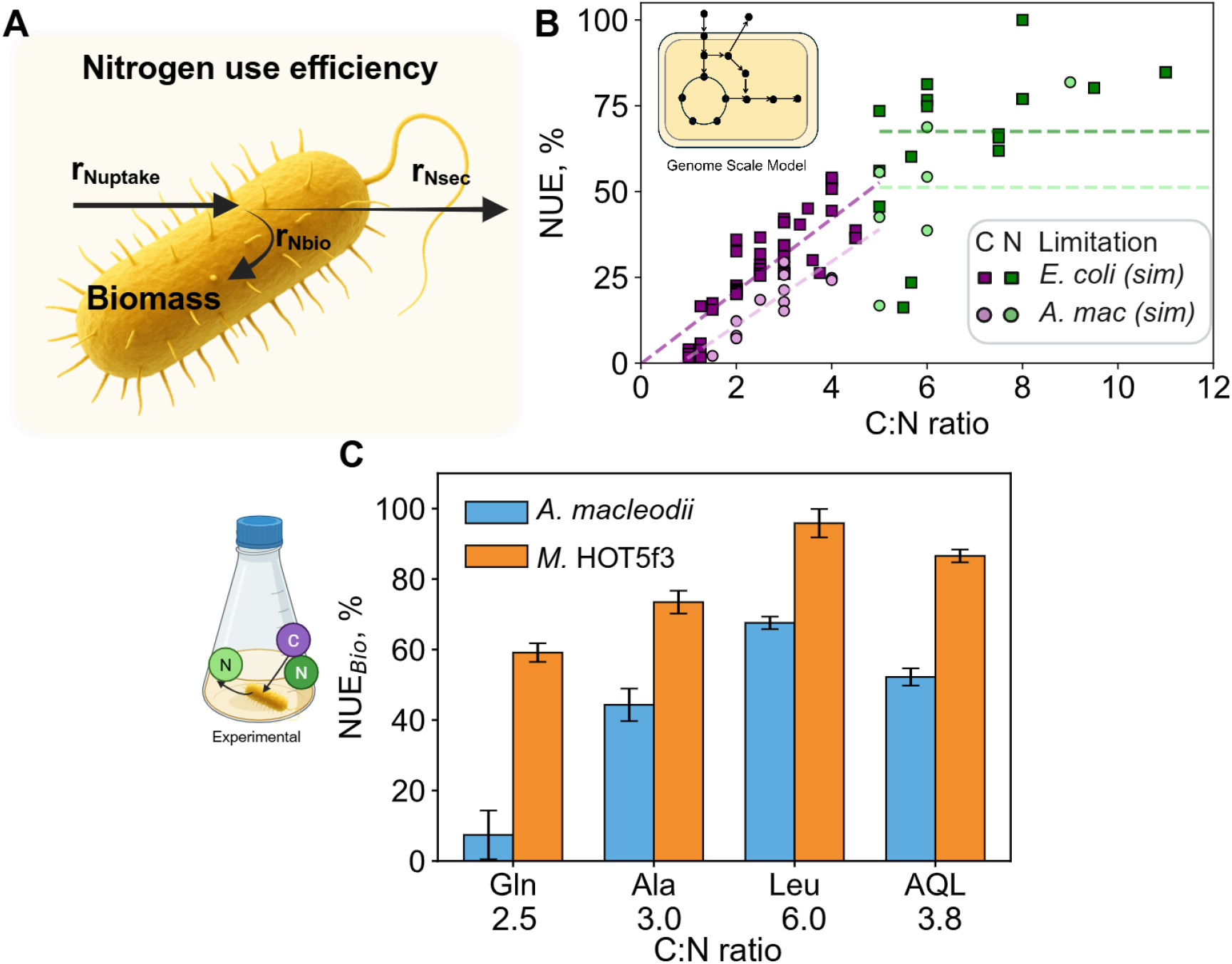
Nitrogen use efficiency across DON substrates from simulations and experiments. **(A)** NUE is defined as the partitioning of nitrogen uptake (*r*_*N*,up_) into biomass incorporation (*r*_*N*,bio_) and secretion (*r*_*N*,sec_). **(B)** FBA-predicted NUE as a function of substrate C:N ratio using the *E. coli* model iML1515 (squares) and the *A. macleodii* model iHS4156 (circles). Under carbon limitation, NUE increases approximately linearly with C:N ratio (*E. coli*: slope = 10.6, *p* < 10^−16^; *A. macleodii* : slope = 9.3, *p* = 1.02 10^−3^), and reaches a plateau under nitrogen limitation (*E. coli*: mean = 67.5%; *A. macleodii*: mean = 51.2%). All nitrogen secretion fluxes are included in the NUE calculation. **(C)** NUEBio for *A. macleodii* MIT1002 and *Marinovum* HOT5f3 grown on alanine (Ala), glutamine (Gln), leucine (Leu), and their mixture (AQL). Ammonium release served as a proxy for *r*_*N*,sec_.

Given the importance of this process, we asked, for some of our strains, whether they exhibit distinct NUE across environmental C:N regimes [19]. Before estimating NUE from experimental data, we applied genome scale modeling to *E. coli* and *A. macleodii* MIT1002 to predict their dependence on substrate C:N ratio during growth on DON (Figure 3**B**). In these simulations, nitrogen uptake and secretion fluxes are tracked, enabling direct quantification of total nitrogen secretion. Consistent with the model of [19], FBA predicts that NUE increases approximately linearly under carbon-limited conditions (C:N 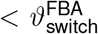) and reaches a plateau once the substrate C:N ratio exceeds 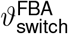. Notably, the two genome-scale models differed quantitatively, with the *A. macleodii* model exhibiting consistently lower NUE values. In several simulations, the *A. macleodii* model predicted allantoin secretion, suggesting a greater loss of nitrogen through excretion and consequently lower NUE.

To assess NUE experimentally, we focused on *A. macleodii* MIT1002 and *Marinovum* HOT5f3, two strains previously co-cultured with *Prochlorococcus* spp. (see Methods) [41, 42, 44]. Both were grown on three amino acids (Gln, Ala, Leu), individually and as a mixture, as sole carbon and nitrogen sources. Nitrogen secretion was quantified by measuring ammonium accumulation in the supernatant over time, which served as a proxy for total nitrogen release. We note that other nitrogen-containing metabolites may also be secreted; however, our analysis focuses on nitrogen remineralization. Consistent with this assumption, FBA simulations predicted ammonium to be the dominant nitrogen-containing secretion product when growth occurred on DON substrates as carbon and nitrogen source. Based on these measurements, we estimated NUE using an alternative formulation (NUE_Bio_) that is more robust to experimental uncertainty and transient non-optimal growth (see Supplementary Figure S14 and Table 8). In analogy with the FBA results, we then plot NUE as a function of substrate C:N ratio (Figure 3**C**). Both strains show increasing NUE with increasing C:N ratio, with highest NUE on leucine. This reflects increasing nitrogen limitation relative to carbon as the substrate C:N ratio rises. Beyond this stoichiometric effect, *Marinovum* HOT5f3 consistently exhibits higher NUE than *A. macleodii* MIT1002, indicating strain-specific metabolic differences. This suggests that under certain conditions *A. macleodii* MIT1002 may display stronger support for surrounding community members that rely on reduced nitrogen sources. Although data on NUE in marine bacteria are scarce, the trends observed here and in FBA simulations are consistent with patterns reported for soil microbial communities [19].

### Does the nitrogen preference depend on the carbon source?

The dataset from experiment E1, conveys how individual simple carbon and nitrogen substrates contribute to growth for a given strain. A natural assumption is that carbon and nitrogen preferences are largely independent, i.e. that the ability of an organism to utilize a particular carbon (nitrogen) source does not depend strongly on the accompanying nitrogen (carbon) source. However, it is not obvious whether this should always be the case, as C and N metabolic pathways are often coupled, at least through the sharing of common cofactor pools. Our dataset gives us the opportunity to revisit this assumption of independence by testing whether specific combinations of carbon and nitrogen substrates are more favorable than others. For each bacterium, the growth yield on all combinations of carbon and nitrogen sources can be represented in the form of a 3×3 matrix (see Figure 4**A**). This provides a visual intuition of the possible interdependence between nitrogen and carbon. In this specific example, nitrate seems to be a good nitrogen source if glucose is the carbon source, but it is not as good if pyruvate is the carbon source (see Figure 4**A**). To quantify the extent of the interdependence between carbon and nitrogen sources across the whole dataset, we used a metric analogous to *epistasis* in genetics, where the effect of one genetic perturbation may depend on the presence of another [43, 45]. In our case, the perturbation is a switch from one substrate to another substrate (e.g., nitrogen source N to N’, or carbon source C to C’). Based on the OD_600_ yield change caused by each of these perturbations, one can estimate the expected additive effect of both perturbations together. Epistasis (*ϵ*) quantifies the deviation from this expectation. The null expectation of independence (*ϵ* = 0) and deviations from it (*ϵ*≠ 0) can be visualized as yield responses to these substrate perturbations (see Figure 4**B**).

**Figure 4:**
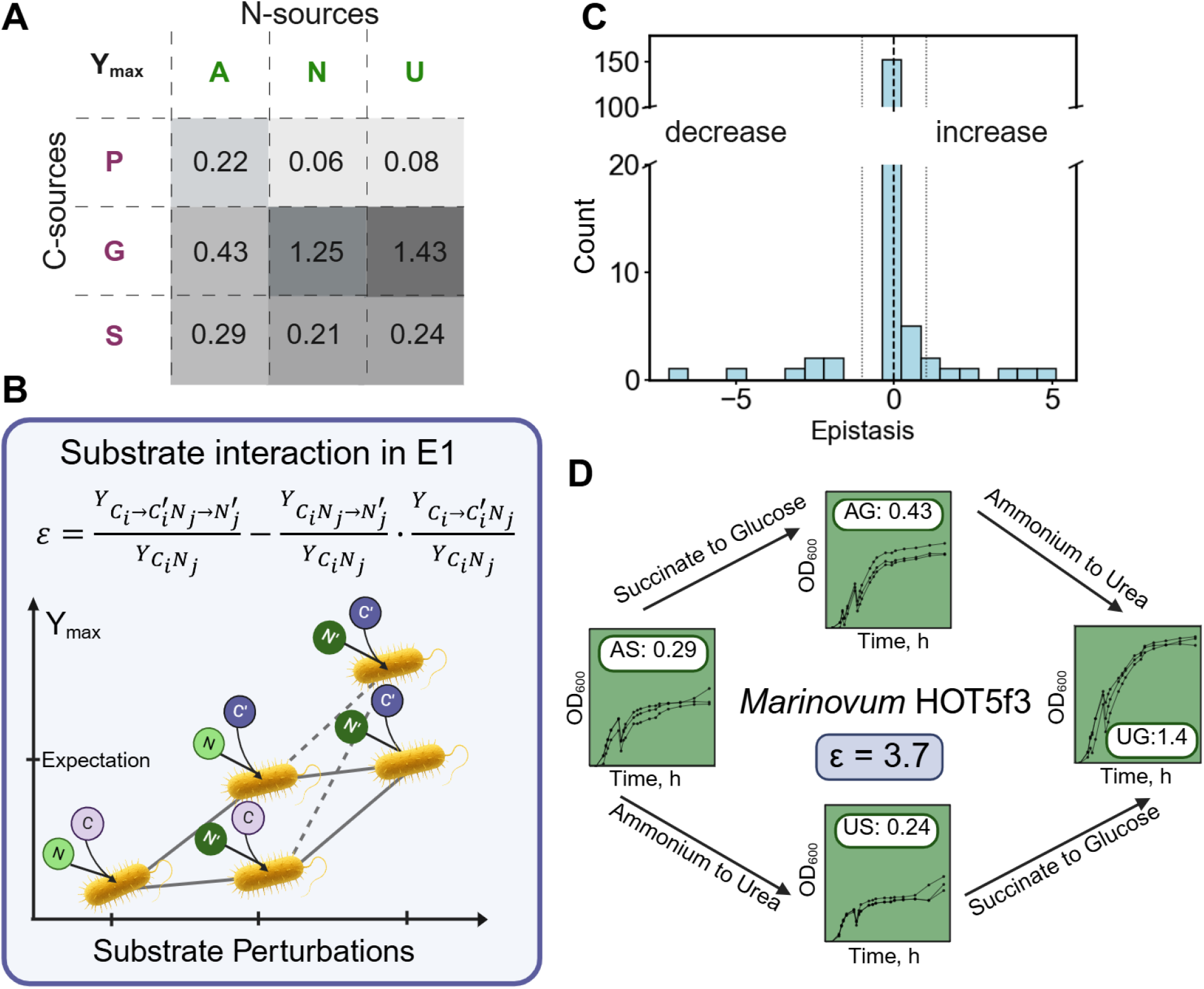
Growth is largely additive with rare epistatic outliers across C and N substrate matrix. **(A)** The DIN dataset (E1) was used to investigate interactions between carbon and nitrogen sources using a metric analogous to *epistasis* in genetics. An example of Y_*max*_ for *Marinovum* HOT5f3 is shown in the 3×3 matrix. Columns represent nitrogen sources (A = ammonium, N = nitrate, U = urea), and rows represent carbon sources (P = pyruvate, G = glucose, S = succinate). **(B)** Schematic representation of the null expectation assuming independent contributions of carbon and nitrogen sources. Deviations from this expectation define epistasis (*ϵ*), with positive values (dashed lines) indicating enhanced and negative values reduced effects. Epistasis equation: 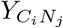 = growth under the reference condition; *Y*_*Ci*→*Ci*_*′,Nj*→*Nj′* = growth when both carbon and nitrogen sources are changed; *Y*_*Nj*→*Nj*_*′* = growth when only nitrogen changes (carbon fixed); *Y*_*Ci*→*Ci*_*′* = growth when only carbon changes (nitrogen fixed). **(C)** Across most strains and substrate combinations, growth reflects independent contributions (*ϵ* ≈ 0, within ±1 SD). Only a few outliers show positive (*ϵ >* 0) or negative (*ϵ* < 0) epistasis. **(D)** Example of positive epistasis (*ϵ* = 3.7) for *Marinovum* HOT5f3 under the conditions AP (ammonium + pyruvate), AG (ammonium + glucose), UG (urea + glucose), and UP (urea + pyruvate).

In our data, the distribution of *ϵ* values across all strains is centered at zero, suggesting that most yields exhibit independent behavior with respect to carbon and nitrogen source combinations (Figure 4**C**). In other words, for most strains, the usability of a given nitrogen (or carbon) source can be described largely independently of the accompanying carbon (or nitrogen) source. However, a small number of strains in specific conditions showed significant deviations from *ϵ* = 0 (7 with *ϵ >* 0 and 7 with *ϵ* < 0). One such case is shown in Figure 4**D** for *Marinovum* HOT5f3. With ammonium and succinate, the strain reached an OD_600_ yield of 0.29 (after subtraction of the negative control). Replacing ammonium with urea slightly decreased the yield, whereas replacing succinate with glucose nearly doubled it. Strikingly, switching both substrates simultaneously to urea and glucose increased the yield to 1.4, corresponding to an epistasis value of *ϵ* = 3.7. This indicates a significant synergistic epistasis effect when *Marinovum* HOT5f3 utilizes urea and glucose together. Despite these individual instances of epistasis, we did not identify any specific pair of carbon–nitrogen sources that was consistently superior across all strains (see Supplementary Figure S15). An important caveat in interpreting these results is that changes in optical density may reflect factors unrelated to growth, such as intracellular storage compounds. In addition, the observed epistatic effects may depend on the limiting nutrient; here, we analyzed epistasis under nitrogen-limiting conditions (see Supplementary Figure S16). Overall, while it is generally valid to state that an organism grows well on a given nitrogen (or carbon) source, for some strains this assessment is context-dependent.

### From amino acids to mixtures to peptides: nonlinearity of nitrogen source complexity

While interactions between simple carbon and nitrogen substrates were generally additive, natural marine environments contain substantially more complex mixtures of organic compounds. We therefore next asked whether increasing molecular complexity in the form of amino acid mixtures (Experiment E4) and peptides (Experiment E5) gives rise to stronger nonlinear growth responses.

To test whether growth followed additive expectations or whether mixtures of multiple C–N compounds generated nonlinear interactions during uptake or intracellular processing, we computed the log_2_-fold change (log(FC)) between the yield of each amino acid mixture and the mean yield of the corresponding individual amino acids (see equation in Figure 5**A**). Under a purely additive scenario, growth on an amino acid mixture would equal the mean yield of the corresponding individual amino acids, resulting in log(FC) = 0 (for normalization see Supplement Text S17, Table 9). The resulting carbon-normalized distribution of log(FC) was approximately bell-shaped, with most conditions across strains clustering around zero, indicating largely additive behavior (see Figure 5**B**). We next analyzed these log(FC) at the level of individual strains to assess how both the magnitude and direction of deviations vary with increasing amino acid mixture size (for normalization see Figure S18). Interestingly, different strains exhibited distinct response patterns (Figure 5**C**). Some strains, such as *Alteromonas macleodii* MIT1002 and *Marinovum* HOT5f3, showed relatively flat responses across increasing mixture size. However, the FC itself varied, suggesting differences between these strains in how they process complex resource mixtures. For *A. macleodii* MIT1002, the values remained largely within one standard deviation of the mean, consistent with an additive response to increasing number of amino acids. In contrast, fold changes for *Marinovum* HOT5f3 were consistently shifted toward negative deviations, meaning that growth on single amino acids had a tendency for higher yields. In a few cases, the log(FC) displays a strong dependence on the number of amino acids. Two extreme examples are *Croceibacter atlanticus*, which exhibited a pronounced increase in yield in mixtures containing more than two amino acids and *Ruegeria pomeroyi* dss3, which showed reduced yields in triple and quadruple mixtures (see Figure 5**C**). Overall, this analysis suggests that, on average, most bacteria can maintain steady growth property at increasingly complex mixtures of amino acids, perhaps reflecting the fact their physiology is dominated by the amount of C and N, irrespective of the source - a sort of amino acid generalism. In contrast, however, some bacteria seem to greatly benefit from diversity of amino acids in the medium. Others, more surprisingly, display an opposite trend, i.e. seem to pay a price for the presence of multiple amino acids, even if none of amino acids is in itself toxic or inhibitory.

**Figure 5:**
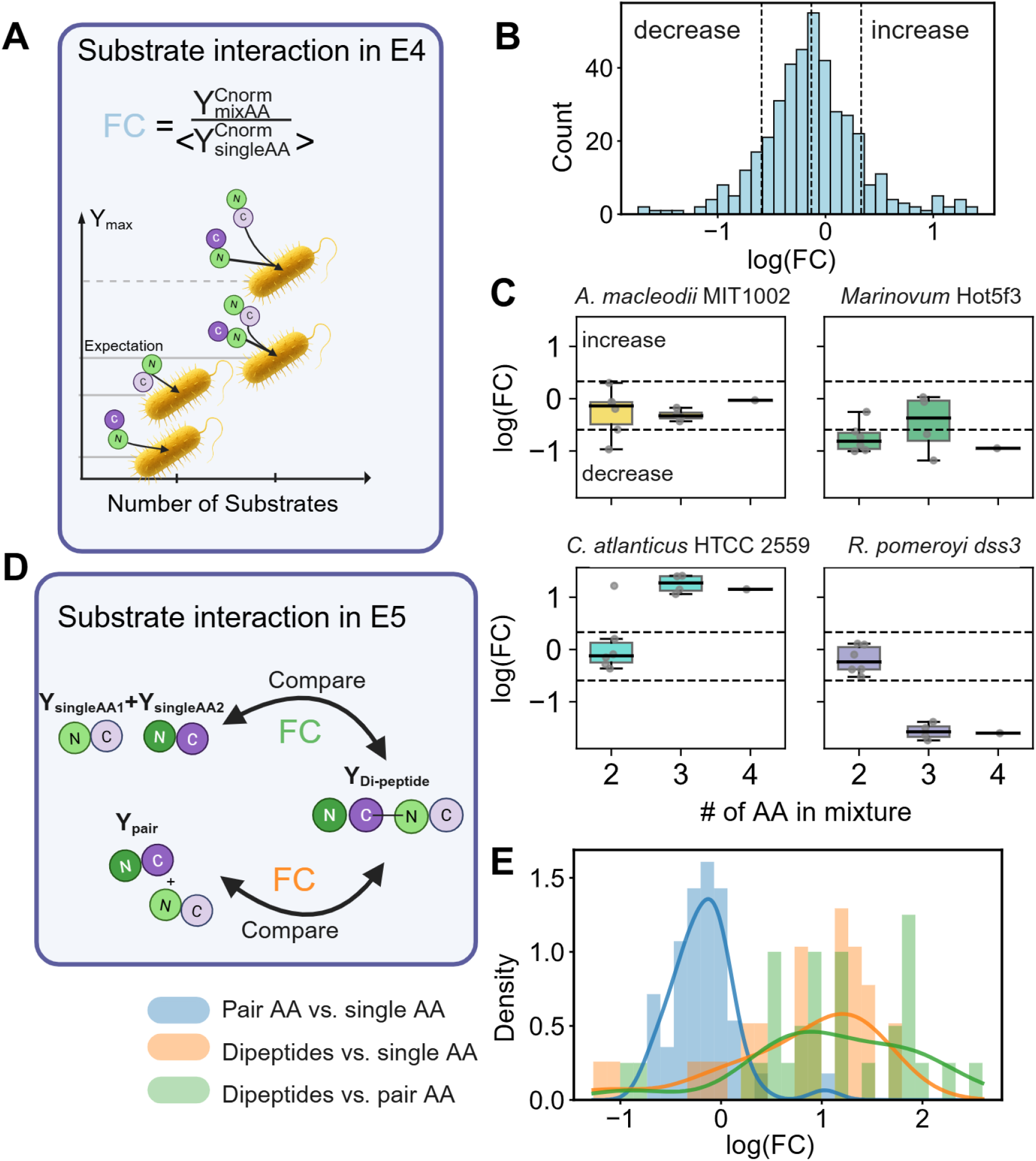
Non-linear growth contributions in amino acid mixtures and dipeptides. **(A)** The null expectation assumes additive contributions from individual amino acids, while deviations indicate nonlinearity (dashed line: positive interaction). **(B)** Log_2_ fold change (log(FC)) comparing the yield of amino acid mixtures (2–4 components) to the mean yield of the corresponding individual amino acids. Values > +1 SD indicate positive interactions, and values < 1 SD indicate negative interactions. **(C)** log(FC) versus size of amino acid mixtures (2, 3 or 4 amino acids) across examples of individual strains, highlighting variability in interaction strength. Dashed lines indicate ±1 standard deviation of the distribution. **(D)** Yields from dipeptides were compared to those from single amino acids (E2) and amino acid pairs (E4) using fold changes. **(E)** Comparisons between single amino acids and amino acid pairs, dipeptides and single amino acids (Distribution in green: 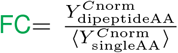), and dipeptides and the corresponding amino acid pair mixtures (Distribution in orange: 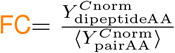)

Growth data on amino acids linked into dipeptides gave us the opportunity to additionally determine whether the bacteria favor dipeptides over the corresponding individual amino acids or amino acid mixtures (Figure 5**D** and Figure 1**A** E5). In particular, we compared the following three quantities: (i) the mean yield of two amino acids supplied individually, (ii) the yield on a pair of amino acids supplied together as a mixture, and (iii) the yield on the corresponding dipeptide, for three of the screened dipeptides (Ala–Gln, Ala–Leu, and Ala–Pro, see Supplement S7). We again used log(FC), as described above, to quantify potential differences. Carbon normalization was performed as before (see Supplement Text S17). Specifically, we compare dipeptide yields to both the mean yield of the individual amino acids and the yield of their mixture. This allows us to test whether covalent coupling of amino acids leads to different growth outcomes compared to supplying the same components separately, either individually or as a mixture Figure 5**D**.

Overall we observed that dipeptides resulted in positive log(FC) relative to both amino acid pairs and the mean yield over individual amino acids. In other words, bacteria tend to prefer a dipeptide rather than the corresponding individual amino acids or their mixture (see Figure 5**E**). These differences suggest strain-specific constraints related to peptide uptake and processing. For example, *E. coli* encodes multiple peptide transporter systems which are described to be more promiscuous than amino acid transporters [46]. Our results are consistent with the abundance of necromass proteins and corresponding proteases in ocean ecosystems, and suggest that either the distribution of peptides or the balance between cost of import and benefit make short peptides a good resource, potentially preferable to individual amino acids [47, 48].

## Discussion

We investigated the growth of diverse marine heterotrophs across a compendium of inorganic and organic nitrogen substrates of increasing molecular complexity. Our analysis revealed broad versatility in nutrient utilization, with metabolic phenotypes largely decoupled from phylogeny, yet strongly structured by substrate properties. In particular, biomass yields can be organized by substrate C:N stoichiometry, degree of reduction, and resource pool complexity, linking nutrient limitation, cellular elemental demands, and energetic constraints.

Our phenotypic data add new knowledge towards the detailed characterization of marine heterotrophic bacteria specifically chosen because of their potential role in modulating primary productivity and their involvement in inter-microbial interactions in the oceans [17, 24, 49, 50]. While some of these organisms had already been studied in detail, and may be effectively considered model organisms of marine heterotrophy, others had been barely characterized before. Thus, our phenotypic landscape can be viewed as a window into a previously unknown aspect of the metabolic diversity of marine bacteria. In addition to being relevant for understanding the lifestyles of bacteria in natural environments, this knowledge could be helpful for designing defined microbial communities.

While many prior phenotyping experiments were focused on the carbon source, our screening is centered on the combined role of carbon and nitrogen in determining bacterial growth. In designing our experiments and analyzing our data we focused our attention on the assimilatory side of nitrogen metabolism. However, we cannot rule out the possibility that nitrate or other compounds could also be involved in dissimilatory metabolism. In the future, it would be interesting to expand the analysis of carbon-nitrogen pairing to conditions where nitrogen may play a more prominent energetic role, e.g. under oxygen limited conditions [51]

The observed relationship between biomass yields and substrate C:N ratio is consistent with a framework previously developed for soil microbial communities (see Supplement Text S11 and Figure S12) [19]. It is notable that a relationship linking nutrient limitation, substrate C:N ratio, and growth efficiency, originally derived for soil communities, also describes the average behavior of diverse marine heterotrophic bacteria across many growth conditions. This suggests that similar quantitative links between substrate stoichiometry, nutrient limitation, and growth yields may operate across microbial habitats. In our framework, yields are linked to *ϑ*_switch_, marking the transition between limitations. This parameter takes into account the carbon lost for respiration and maintenance, and thus represents an upper bound for the biomass C:N ratio.

The efficiency with which biomass is produced from a substrate depends not only on its stoichiometry but also on its energetic content, quantifiable through the degree of reduction of the carbon. This dependency, which has its roots in classical biological thermodynamics [52, 53] has been recently revisited in the broader context of microbial ecology, suggesting that substrate properties are a fundamental determinant of ecosystem microbial physiology [30].

Our analysis sheds some new light on the relationship between DON usage and ammonium exudation and exchange. We suggest that NUE may be used as a valuable community-relevant ecological trait, capturing the capacity of some bacteria to recycle nitrogen for the benefit of other organisms. Importantly, our experiments quantified ammonium as the only exudate. A full understanding of the relationship between NUE and specific nitrogen exudates will require complete nitrogen balances based on comprehensive flux, biomass and exudate measurements. Such experiments could enable systematic, time-resolved estimates of NUE along the growth curve, providing insight into metabolic strategies underlying nitrogen loss. Further work could also explore in more depth some aspects of nitrogen metabolism that cannot be easily implemented in a high throughput manner, including the potential role of air-mediated ammonia cross-feeding [26] and the relevance of dual limitation regimes [54, 55].

We envisage several directions for future work building on our results. First, metabolic capabilities are likely to depend strongly on substrate concentrations and environmental variables such as temperature, pH, and salinity. Parameters relevant to inter-species interactions, including NUE, may therefore vary substantially across environmental contexts and should be explored in depth experimentally [56, 57]. It would also be interesting to extend microbial phenotyping to larger sets of strains, substrates, and nutrient combinations, including additional elements such as phosphate. Furthermore, future work could incorporate community and spatial aspects of microbial metabolism, as aggregation or particle colonization are likely closely linked to nutrient utilization strategies [16, 58]. Ultimately, systematic mapping of microbial metabolic landscapes across environmental space may help refine predictive and quantitative understanding of how the lifestyles of different bacteria shape biogeochemical processes [59–61].

## MATERIALS AND METHODS

### Experimental Methods

#### Media and strain handling for screenings

Strains were streaked from cryo stocks onto marine broth agar plates (Difco 2216, 1.5% agar) and incubated for 5–7 days at 26 °C. For screening, single colonies were inoculated into 3 mL liquid marine broth (Difco 2216) and grown overnight at 26 °C and 200 rpm in a Thermo Scientific MaxQ 6000 incubator. Pre-cultures were then transferred (1:10 v/v) into freshly prepared nitrogen-limited medium. The base medium (see Supplement Table 1 and [17]) was supplemented with an N-solution (final concentrations: 2.02 mg L^−1^ KNO_3_, 1.1 mg L^−1^ NH_4_Cl) and a C-solution (1 g L^−1^ each of glucose and pyruvate). All screenings were performed at 26 °C without shaking in 96-well plates (200 µL volume). Plates were mixed prior to optical density measurements at 600 nm, which were taken at least once per day. Carbon and nitrogen sources and concentrations are listed in Table 1. For some conditions, the medium was adjusted to pH 7.4 with NaOH.

**Table 1:** Concentration of carbon and nitrogen in screenings.

| Screening | C conc., mM | N conc., mM |
| --- | --- | --- |
| DIN: | 100 | 5 |
| Amino acid | 100<sup>1</sup> | 7.8 |
| Polyamine | 100<sup>1</sup> | 7.8<sup>2</sup> (6.9<sup>2</sup>) |
| Nucleotide/Nucleoside | 100<sup>1</sup> | 7.8 |
| Amino acid Mixture | no carbon | 10 |
| Dipeptide | 100<sup>1</sup> | 10 |

<sup>1</sup> If carbon was added, it was a 1:1 mixture of glucose and pyruvate.
<sup>2</sup> Spermidine (spermine)

#### Strain handling for N-oligotrophy

To assess nitrogen oligotrophy, heterotrophic strains were first plated on marine broth agar (Difco 2216, 1.5 %), then transferred to minimal medium plates, followed by two transfers onto nitrogen-starvation plates lacking added nitrogen. Minimal medium plates were prepared by mixing 2× liquid base medium with 2× H_2_O-agar (1.5% w/v) (see Supplement Table 1 and [17]) and supplemented with 2 g L^−1^ each of pyruvate and glucose, 0.26 g L^−1^ NH_4_Cl, and 0.537 g L^−1^ KNO_3_. Nitrogen-starvation plates were prepared identically but without nitrogen. Plates were incubated at 26 °C for one week before growth was assessed.

#### NUE shake flask experiments

Pre-cultures were prepared as described above. After nitrogen starvation, 0.5 mL of the second pre-culture was inoculated into 24.5 mL of base medium containing 10 mM nitrogen supplied as the respective amino acid or a mixture. Cultures were grown at 26 °C with shaking at 200 rpm for at least five days. Samples were collected, filtered, and used for enzymatic ammonia quantification (Megazyme ammonia assay).

### Computational Methods

#### Inference of growth parameters

OD_600_ measurements were normalized to the initial value and log-transformed. Growth was determined using a threshold derived from negative controls. Growing strains were fitted with a Gompertz function to estimate µ_max_, *Lag* phase, and Y_max_. For non-growers, µ_max_ was set to 0, lag phase to 50 h, and Y_max_ was obtained directly from the data. Statistical comparisons were performed against growing negative controls (see Supplement Figure S8).

#### Principal component analysis

Growth parameters obtained from experiments E1, E2, and E5 were filtered based on *Y*_*max*_, standardized using the StandardScaler implementation in scikit-learn, and subjected to principal component analysis (PCA). In the strain-based PCA, data points were colored according to phylogenetic class. To investigate patterns across substrates, the transposed data matrix was normalized and analyzed by PCA. Data points were colored according to substrate class (Supplementary Figure S9).

#### Flux balance analysis simulation

Standard FBA (COBRApy) was performed using the genome-scale metabolic model of *A. macleodii* MIT1002 (iHS4156 v3.0.0) and the *E. coli* model iML1515 [62]. Simulations were used to predict growth rates on different DON substrates provided as sole carbon and nitrogen sources. Nitrogen uptake was constrained to 10 mmol N gCDW^−1^ h^−1^, and oxygen uptake was limited to 15 mmol gCDW^−1^ h^−1^.

#### Yields, Degree of reduction and NUE

For the FBA simulations, uptake rates of the respective DON substrates were used to calculate *Y*_*N*_ and *Y*_*C*_. For the experimental data, we assumed that all supplied substrate was consumed by the end of the experiment, allowing us to approximate uptake using the total amount of carbon and nitrogen supplied. Here, *N*_*total*_ and *C*_*total*_ denote the total amounts of nitrogen and carbon provided by each substrate.

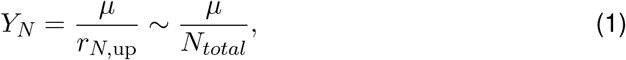

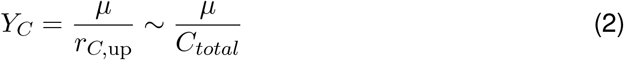

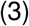

The degree of reduction of carbon (*γ*_*C*_) and nitrogen (*γ*_*N*_) was calculated from the elemental composition of each substrate following [30]:

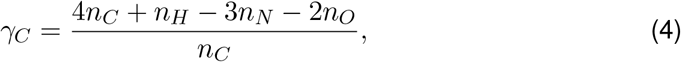

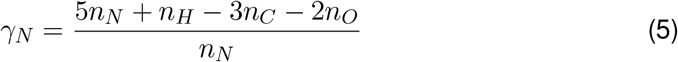

where *n*_*C*_, *n*_*H*_, *n*_*N*_, and *n*_*O*_ denote the number of carbon, hydrogen, nitrogen, and oxygen atoms in the substrate molecule, respectively. *γ*_*C*_ and *γ*_*N*_ therefore represent the average degree of reduction per carbon and nitrogen atom. We used data from experiments E2 and E5 for these analyses

Nitrogen use efficiency was estimated from experimental growth and ammonium secretion measurements as:

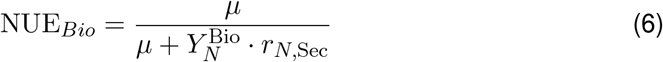

where *µ* is the specific growth rate, 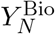 is the biomass yield per unit nitrogen incorporated into biomass, estimated from the *Alteromonas macleodii* MIT1002 genome-scale metabolic model (iHS4156), and *r*_*N*,Sec_ is the nitrogen secretion rate measured as ammonium production (for details see Supplement S13 and S14).

#### Estimates of nonlinearities

OD_600_ values above the growth threshold were used for epistasis calculations:

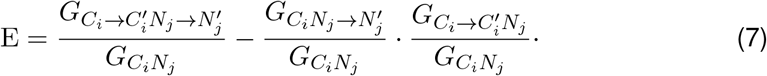

Here, 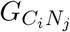 denotes growth on the initial carbon-nitrogen combination, 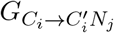 growth after changing the carbon source, 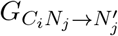 growth after changing the nitrogen source, and 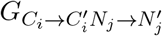 growth after changing both sources simultaneously (see Supplements S15 and S16). To compare growth on dipeptides and amino acid mixtures with growth on the corresponding individual amino acids, log fold changes [log(FC)] were calculated (Supplement S17).

## Acknowledgements

The authors are grateful to Mary-Ann Moran, Rogier Braakman and members of the Segrè, the Sher lab and the c-CoMP community for helpful feedback and comments on the results and the manuscript.

## Data availability

The data underlying this article are available Github.

## Author contributions statement

FK, DSe and DSh designed the study. FK planned and conducted the experiments and the statistical analysis of data. SJ, KH and MO carried specific experiments and assisted with overall experimental setup. HS provided the *Alteromonas* genome scale model. FK and DSe developed mathematical models and computer simulations. FK, DSe and DSh wrote the manuscript, with feedback and contributions by all authors.

