## Supplement for "Organizing principles in the nitrogen–carbon landscape of marine heterotrophic bacteria"

### S0 Additional Material and Methods

Table 1: Marine base medium composition based on [1].

| Compound | Concentration (g $L^{1}$) |
|---|---|
| NaCl | 19.45 |
| $MgSO_4 \cdot 7H_2O$ | 3.24 |
| $MgCl_2$ | 5.90 |
| KCl | 0.55 |
| $CaCl_2$ | 1.80 |
| Boric acid | 0.022 |
| $FeCl_3 \cdot 6H_2O$ | 0.11 |
| $Na_2PO_4$ | 0.008 |
| $NaHCO_3$ | 0.16 |
| Thiamin-HCl | 0.0004 |
| Biotin | 0.000002 |
| B12 (cyanocobalamin) | 0.000002 |
| Folic acid | 0.000004 |
| PABA | 0.00002 |
| Nicotinic acid (niacin) | 0.0002 |
| Inositol | 0.002 |
| Ca Pantothenate | 0.0004 |
| Pyridoxine-HCl | 0.0002 |
| $ZnSO_4 \cdot 7H_2O$ | 0.008 |
| $CoCl_2 \cdot 6H_2O$ | 0.005 |
| $MnCl_2 \cdot 4H_2O$ | 0.09 |
| $Na_2MoO_4 \cdot 2H_2O$ | 0.003 |
| $Na_2SeO_3$ | 0.01 |
| $NiCl_2 \cdot 6H_2O$ | 0.01 |

**Nitrogen starvation plates:** Growth was observed for several strains in media containing only a carbon source and no added nitrogen. To further investigate this phenotype, all strains were streaked onto nitrogen-free marine agar plates (see Methods). After colony formation, single colonies were transferred onto fresh nitrogen-free plates to minimize the possibility that growth was supported by intracellular nitrogen reserves or nitrogen released from dead cells. Plates were incubated for two weeks, after which colony formation was documented photographically (Github, Supplement Table 6). Growth on nitrogen-free plates was compared to growth observed in the DIN screening (E1). Notably, three strains that did not grow under any carbon–nitrogen combination tested in the DIN screen were nevertheless able to form colonies on nitrogen-free marine plates (Supplement Table 2).

**Reannotation:** Genome reannotation was performed using Prokka v1.14.5 with default settings [2]. The genomes used in this study are available at Zenodo.

Table 2: Comparison of non-grower strains from the DIN screening (E1) with growth on nitrogen-starvation plates and with data from [1]. *results from this study; HMBorg = organic acids mixture [1]; HMBntrl = neutral sugars mixture [1].

| Non-grower Strain | Growth DIN Plates* | Yield HMBorg | Yield HMBntrl |
|---|---|---|---|
| Atlcs | 0 | 0 | 0 |
| Exig | 0 | 0 | 0 |
| Fang | 0 | 0 | 0 |
| Kordia | 0 | 0 | 0 |
| Koree | 0 | 0 | 0 |
| Phalo | Growth | 0.12 | 0 |
| Sflavi | Growth | 0.12 | 0 |
| T2 | Growth | 0 | 0 |
| Vcamp | Growth | 0.20 | 0 |
| Sheweden | 0 | 0 | 0 |
| Relong | 0 | 0.15 | 0 |

### S1 DIN Screening - Grower and Non-Growers

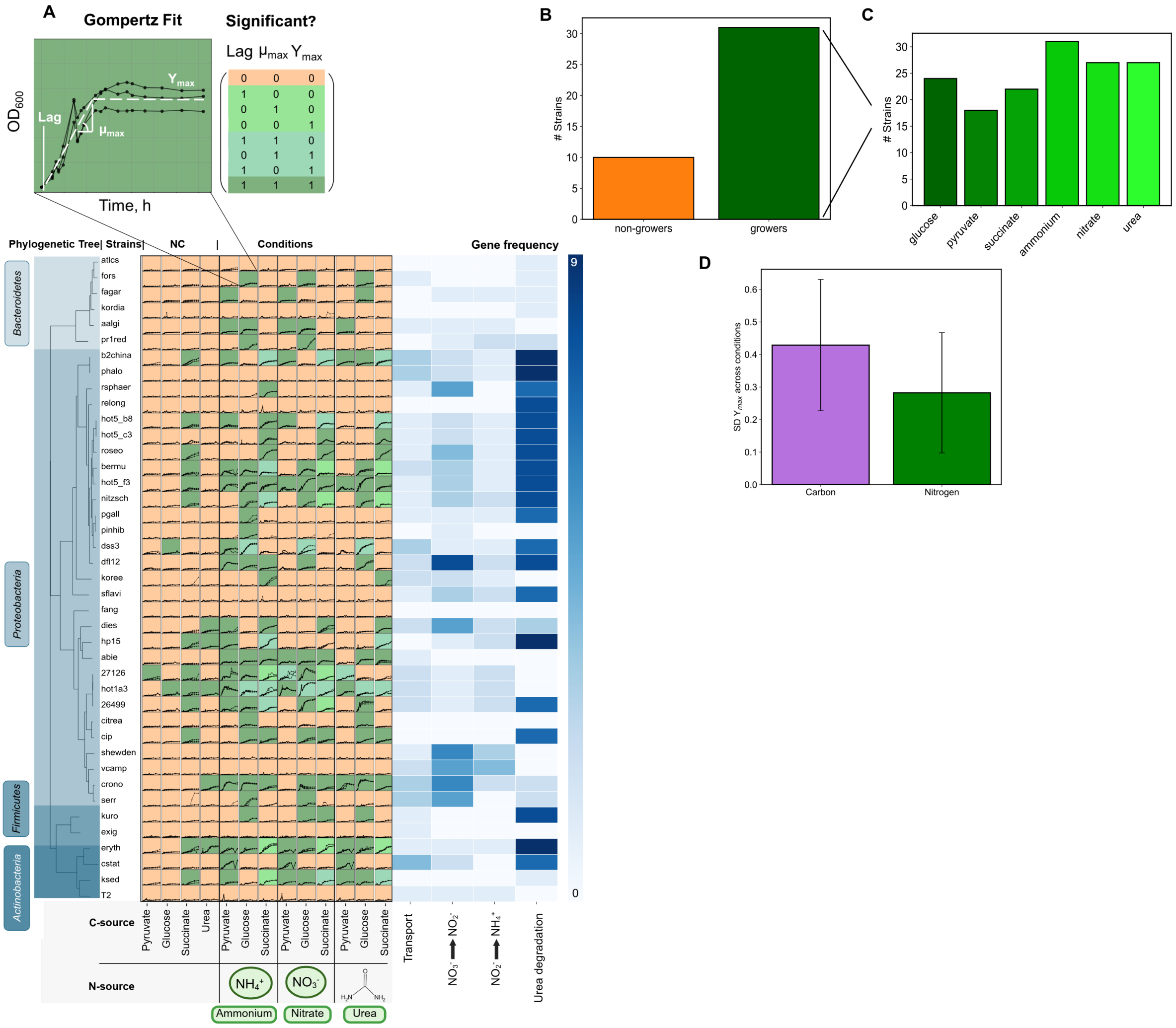


Figure S1: **(A)** Growth curves of marine strains grown on combinations of glucose, pyruvate, and succinate as carbon sources, and ammonium, nitrate, and urea as nitrogen sources. Colors indicate the number of parameters that were significant after Gompertz fitting and comparison with growing negative controls. **(B)** Number of growers and non-growers strains in the DIN screening (E1). **(C)** Number of grower strains growing on different substrates used. **(D)** Variation in Yield ($Y_{max}$) across nitrogen or carbon conditions.

#### S2 nitrate/nitrite and urea transport, reduction and degradation

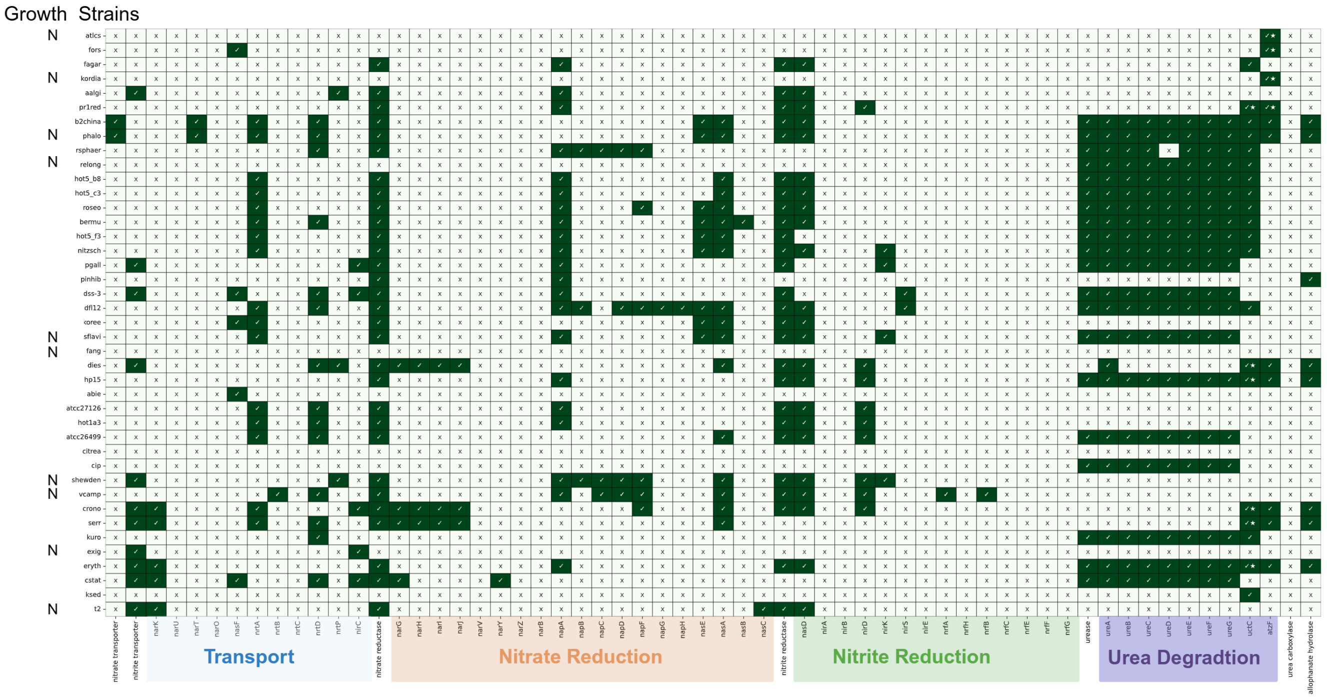


Figure S2: Genes for nitrate/nitrite transport, nitrate reduction, nitrite reduction and urea degradation. The Growth column indicates strains that were unable to grow under DIN conditions. “Nitrate transporter,” “nitrite transporter,” “nitrate reductase,” “nitrite reductase,” “urease”, “urea carboxylase,” and “allophanate hydrolase” are summary columns. A check mark indicates that a gene or related product was found in the annotation. A small star marks annotations identified in other databases, such as NCBI.

Table 3: Genes used as queries to contextualize phenotypic and genomic data related to nitrate/nitrite transport and reduction.

| Genes | Product / Reaction | Note |
|---|---|---|
| Transport | | |
| *nasD, nasE, nasF* | Nitrate ($NO_3^-$) transport | ABC-type transporter; ATP-dependent |
| *nrtA, nrtB, nrtC* | Nitrate ($NO_3^-$) transport | Nomenclature derived from cyanobacteria (e.g. *Synechococcus elongatus* PCC 7942) |
| *narK, narU, narT* | Putative nitrate/nitrite transporter | Major facilitator superfamily-type transporters |
| *nrtA, nrtB, nrtC, nrtD, nrtP* | Nitrate/nitrite transporter | Expanded transporter gene set |
| *nirC* | Nitrite transporter | Inner membrane transporter |
| Nitrate Reduction | | |
| *nasA, nasB, nasC* | Assimilatory nitrate reductase | Cytoplasmic reduction of $NO_3^-$ to $NO_2^-$ |
| *napA, napB, napC, napD, napF, napG, napH* | Periplasmic nitrate reductase | Includes accessory and maturation proteins |
| *narG, narH, narI, narJ, narV, narY, narZ, narB* | Respiratory nitrate reductase | Membrane-bound complex and accessory proteins |
| Nitrite Reduction | | |
| *nasD* | Nitrite reductase [NAD(P)H] | Assimilatory nitrite reductase |
| *nirD* | Nitrite reductase (NADH), small subunit | Cytoplasmic enzyme component |
| *nirK* | Copper-containing nitrite reductase | Denitrification pathway |
| *nirS* | Cytochrome cd1 nitrite reductase | Denitrification pathway |
| *nrfA, nrfH, nrfB, nrfC, nrfE, nrfF, nrfG* | Cytochrome c nitrite reductase | Dissimilatory nitrite reduction to ammonium |

Table 4: Genes used as queries to contextualize phenotypic and genomic data related to urea degradation.

| **Genes** | **Product / Reaction** | **Note** |
|---|---|---|
| Urease genes | | |
| *ureA, ureB* | Small catalytic subunits | |
| *ureC* | Lage, catalytic subunit | widely found in various marine environments [3] |
| *ureD, ureE, ureF, ureG* | Accessory proteins | |
| Amidolyase pathway genes | | |
| *uctC* | urea carboxylase | Transfers a carboxyl group from bicarbonate to urea [4] |
| *atzf* | allophanate hydrolase | Hydrolytic breakdown of allophanate [4] |

Table 5: Genes used as queries to contextualize phenotypic and genomic data related to pyruvate assimilation and transport.

| **Genes** | **Product / Reaction** | **Note** |
|---|---|---|
| Pyruvate assimilation | | |
| *aceE (pdhA), aceF (pdhB), lpd (pdhD)* | Pyruvate dehydrogenase complex | Converts pyruvate to acetyl-CoA, linking glycolysis to the TCA cycle |
| *poxB* | Pyruvate oxidase | Converts pyruvate to acetate; often active under aerobic conditions |
| Pyruvate transporters | | |
| *btsT* | High-affinity pyruvate transporter | Induced under low pyruvate concentrations |
| *cstA* | Carbon starvation protein A (peptide/pyruvate transporter) | Associated with nutrient limitation and alternative carbon uptake |
| TRAP / $C_4$-dicarboxylate transport systems | | |
| *dctP, dctQ, dctM* | TRAP transporter ($C_4$-dicarboxylates) | Involved in uptake of organic acids; may contribute to pyruvate-related metabolism |

### S3 Pyruvate related Genes

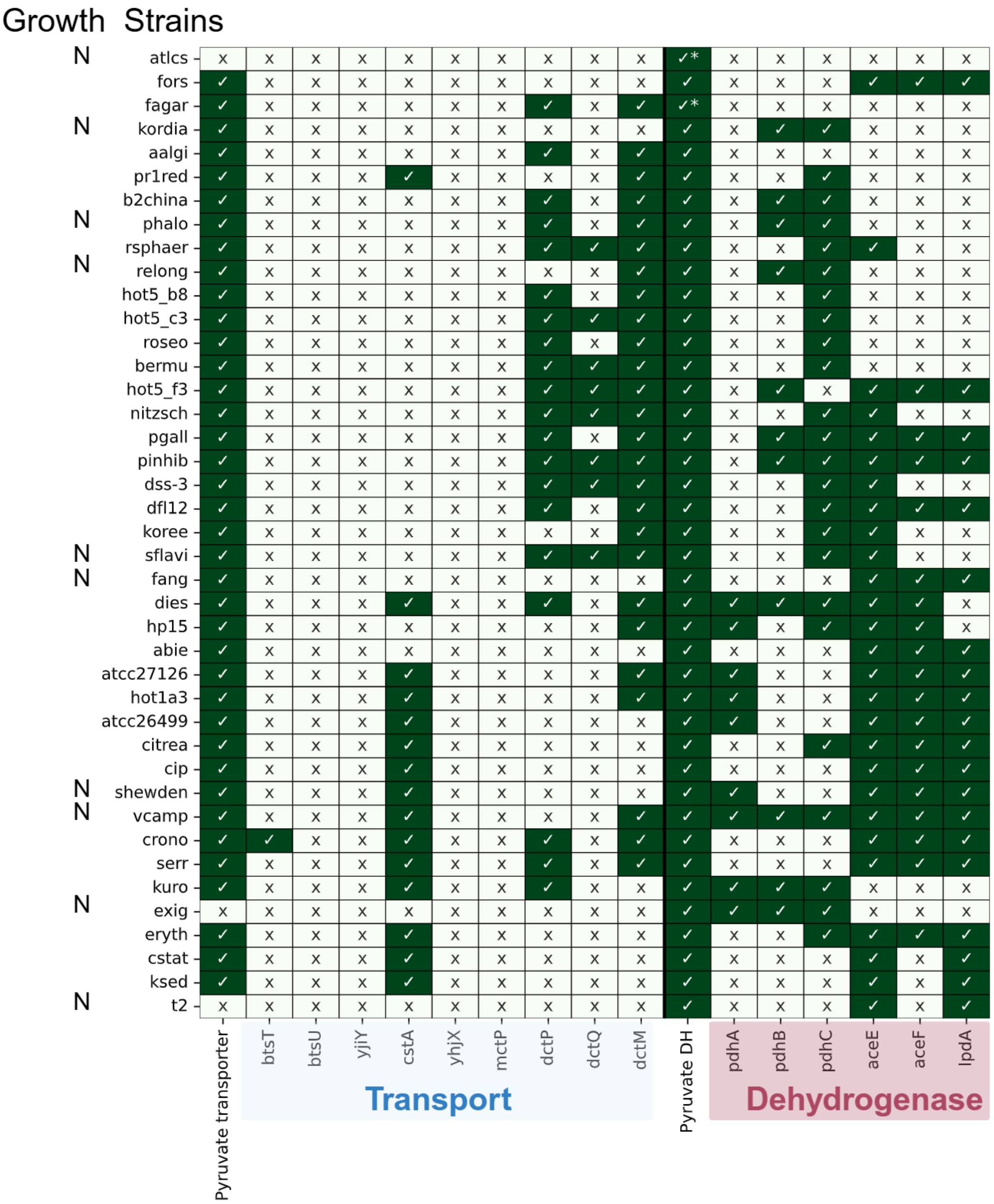


Figure S3: Annotated genes related to pyruvate transport and pyruvate dehydrogenase are shown. The “Growth” column indicates strains unable to grow under DIN conditions (N=non-growers). “Pyruvate transporter” and “Pyruvate DH” are summary columns; a green check mark indicates the presence of any annotated gene or product related to these functions (not all are shown individually). Asterisks denote annotations identified in external databases, including pyruvate dehydrogenases for atlcs (*Croceibacter atlanticus*) GenBank: EAP88180.1 and fagar (*Formosa agariphila*) GenBank: CDF78015.1).

Table 6: Growth (yes ✓/no X) on marine broth (MB), defined medium (MPB), and nitrogen starvation plates (MPB-N; first and second exposure). Not determined values are shown in white.

| Strain_ID | MB? | MPB? | MPB-N? | MPB-N 2nd? |
|---|---|---|---|---|
| 26499 | ✓ | ✓ | ✓ | ✓ |
| 27126 | ✓ | ✓ | ✓ | ✓ |
| Aalgi | ✓ | ✓ | ✓ | ✓ |
| abie | ✓ | ✓ | ✓ | × |
| atlcs | ✓ | × | nd | nd |
| B2china | ✓ | ✓ | × | nd |
| bermu | ✓ | ✓ | × | nd |
| CIP | ✓ | × | nd | nd |
| citrea | ✓ | ✓ | ✓ | nd |
| crono | ✓ | ✓ | ✓ | ✓ |
| cstat | ✓ | ✓ | ✓ | ✓ |
| DFL-12 | ✓ | ✓ | × | nd |
| dies | ✓ | ✓ | ✓ | × |
| DSS-3 | ✓ | ✓ | × | nd |
| eryth | ✓ | ✓ | ✓ | ✓ |
| exig | ✓ | × | nd | nd |
| Fagar | ✓ | ✓ | ✓ | nd |
| fang | ✓ | × | nd | nd |
| fors | ✓ | ✓ | ✓ | × |
| HOT1A3 | ✓ | ✓ | ✓ | ✓ |
| HOT5_C3 | ✓ | ✓ | × | nd |
| HOT5_B8 | ✓ | ✓ | × | nd |
| HOT5_F3 | ✓ | ✓ | × | nd |
| HP15 | ✓ | ✓ | ✓ | × |
| kordia | ✓ | × | nd | nd |
| koree | ✓ | × | nd | nd |
| Ksed | ✓ | ✓ | × | nd |
| kuro | ✓ | ✓ | ✓ | ✓ |
| nitzsch | ✓ | ✓ | ✓ | × |
| Pgall | ✓ | ✓ | ✓ | × |
| PR1red | ✓ | ✓ | ✓ | ✓ |
| Phalo | ✓ | ✓ | ✓ | × |
| Pinhib | ✓ | ✓ | × | nd |
| roseo | ✓ | ✓ | ✓ | × |
| relong | ✓ | × | nd | nd |
| Rsphaer | ✓ | ✓ | × | nd |
| serr | ✓ | ✓ | × | nd |
| Sflavi | ✓ | ✓ | ✓ | × |
| Shewden | ✓ | × | nd | nd |
| T2 | ✓ | ✓ | ✓ | × |
| vcamp | ✓ | ✓ | ✓ | ✓ |
| Mit1002 | ✓ | ✓ | ✓ | ✓ |

#### S4 Growth on Dissolved Organic Nitrogen - DON

Growth on amino acids as sole carbon–nitrogen sources was tested, as well as after supplementation with either ammonium (additional nitrogen = "+N") or a pyruvate–glucose mixture (additional carbon = "+C"). Growth was determined as described in S1 (see Figure S4**A B**). Growth on amino acids alone was common (Figure S4**C**), although in some cases the signal-to-noise ratio was low. To assess whether growth was carbon- or nitrogen-limited, we compared growth on single amino acids with growth after carbon or nitrogen supplementation (Figure S4**C**). Responses not significantly different from growth on the amino acid alone are shown in gray. A significant increase, defined as at least one parameter ($Lag$, $\mu_{\mathrm{max}}$, or $Y_{\mathrm{max}}$) improving relative to the amino acid alone, is shown in green for nitrogen addition and in purple for carbon addition. Overall, growth increases were more frequently observed upon carbon addition, suggesting that growth on amino acids was primarily carbon-limited under these conditions. No clear phylogenetic clustering of growth phenotypes was observed across the tested amino acids (Figure S4**D**).

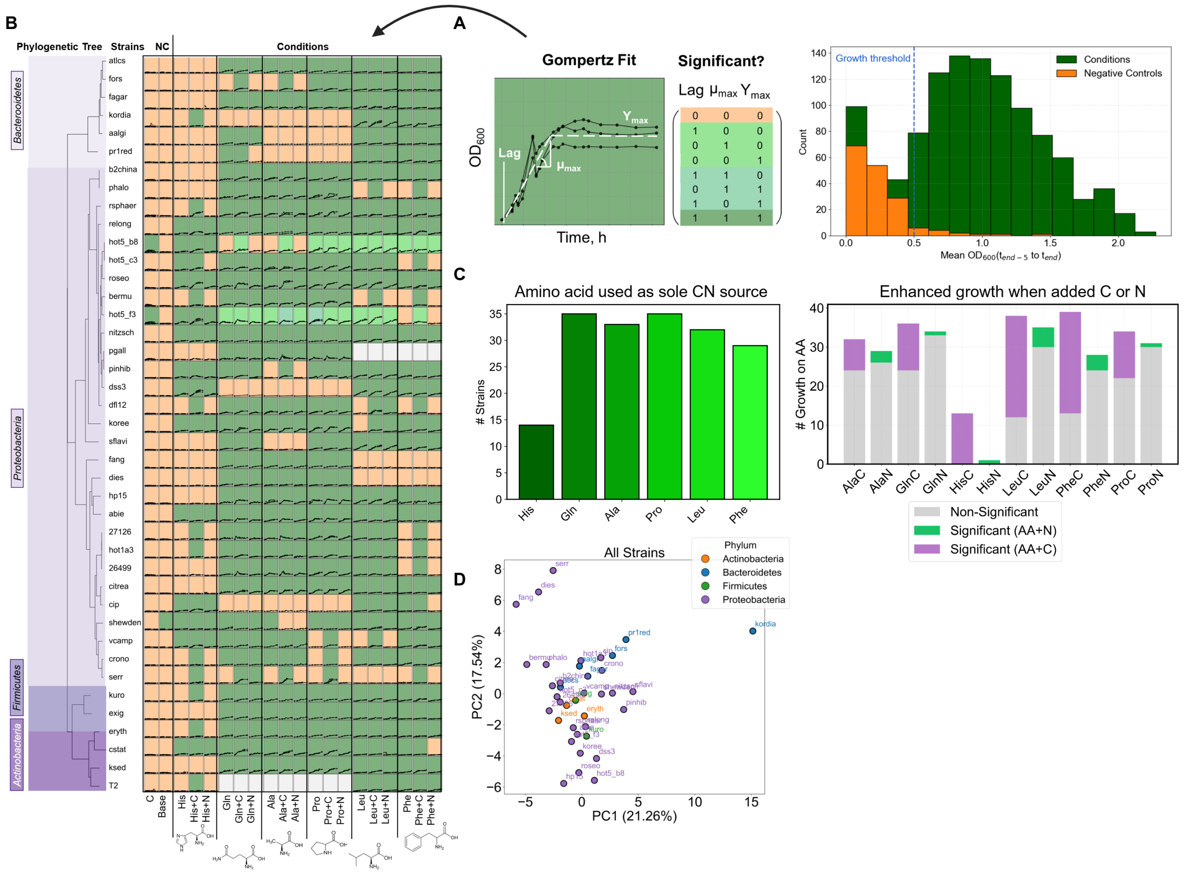


Figure S4: **(A)** Strains exceeding the growth threshold were used for Gompertz curve fitting. The raw growth data are shown in the heatmap in **(B)**. NC = negative control. C-only = carbon mixture provided without nitrogen. Base = no carbon or nitrogen provided. Colors represent the number of significant parameters compared to the respective growing negative controls. **(C)** Number of strains capable of using amino acids as sole carbon–nitrogen sources, and number of strains showing increased growth (defined as at least one parameter, lag, $\mu_{max}$, or $Y_{max}$ being significantly improved relative to growth on the amino acid alone) upon addition of an external carbon source (pyruvate–glucose mixture) or nitrogen source (ammonium). Ala = alanine, Gln = glutamine, Leu = leucine, Pro = proline. **(D)** Principal component analysis (PCA) of growth across all amino acid conditions. Strains are colored according to their phylum.

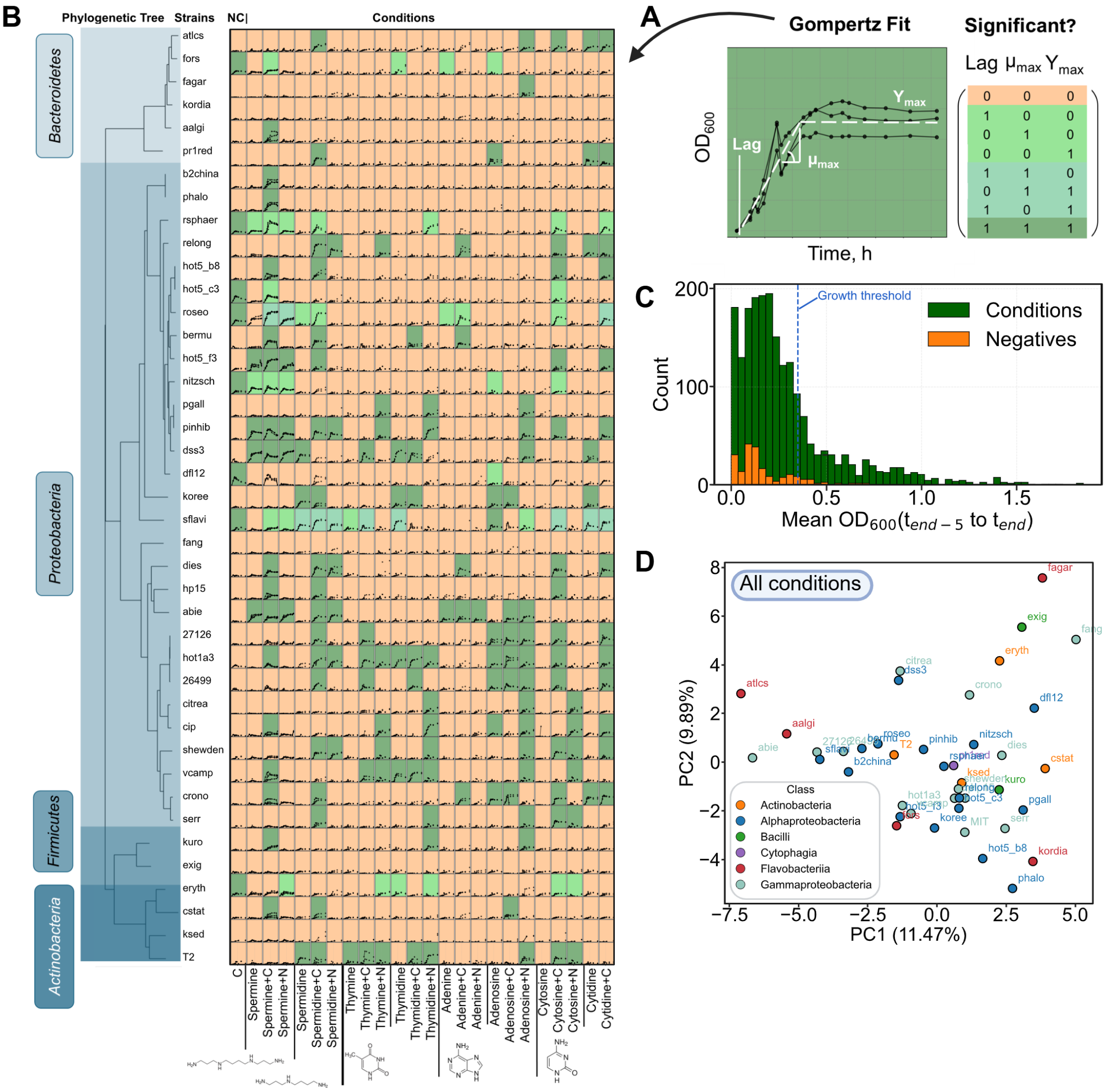


Figure S5: Marine heterotroph growth on polyamines, nucleobases, and nucleosides. **(A)** Colors indicate the number of parameters that are significant after Gompertz fitting and comparison with growing negative controls. **(B)** Heatmap showing growth on the polyamines spermine and spermidine, as well as on nucleobases and their corresponding nucleosides, supplemented with carbon and nitrogen. Note that cytosine was supplemented with carbon only. **(C)** Yield data for defining growth threshold of $\ln(OD_{600})$ = 0.35. **(D)** PCA of all conditions; colors indicate heterotroph classes.

#### S6 Growth of marine heterotrophs on different amino acid mixtures

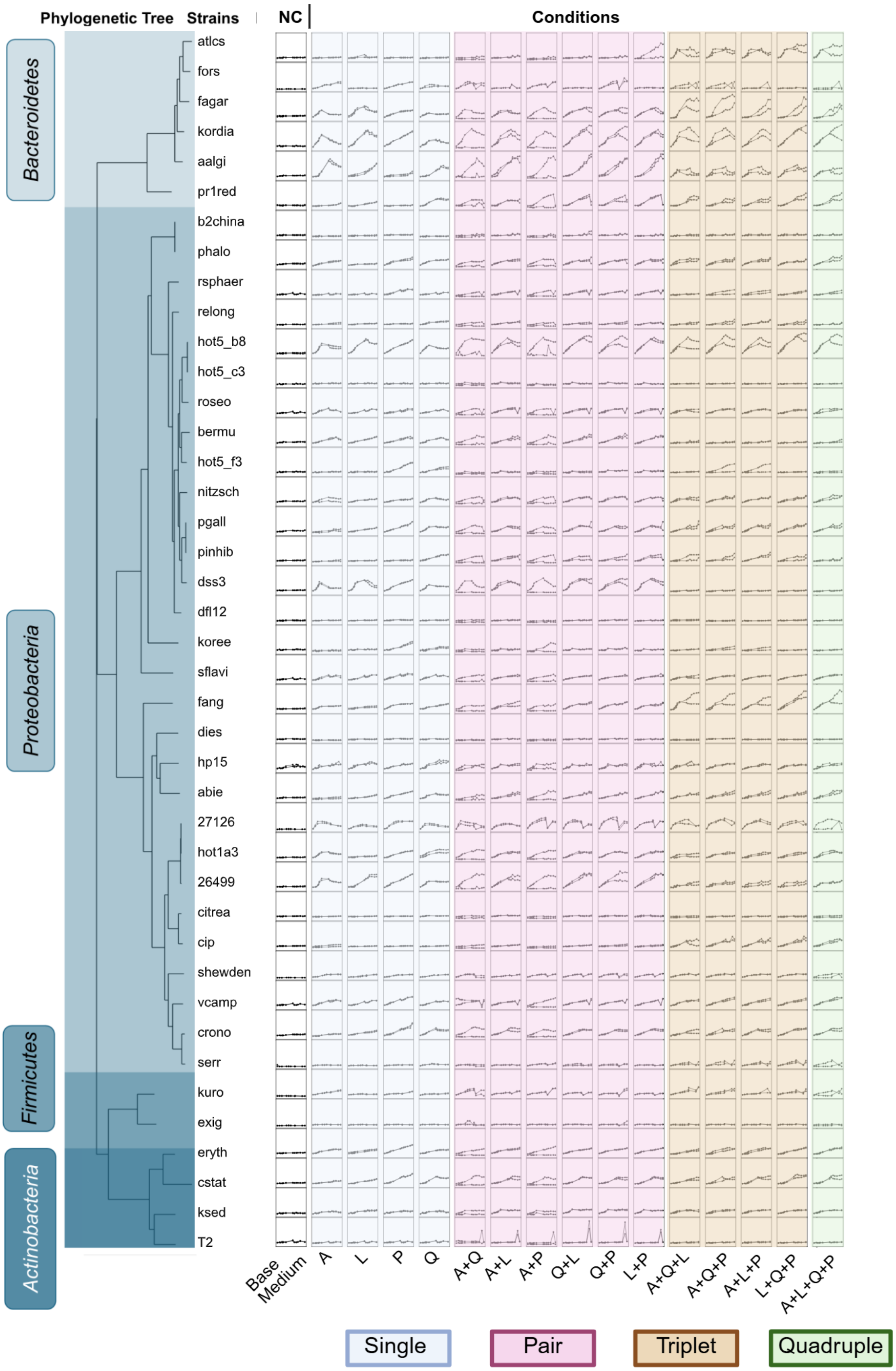


Figure S6: Growth on individual amino acids and mixtures of pairs, triplets, and quadruplets. All experiments were performed in duplicates.

### S7 Growth on dipeptides

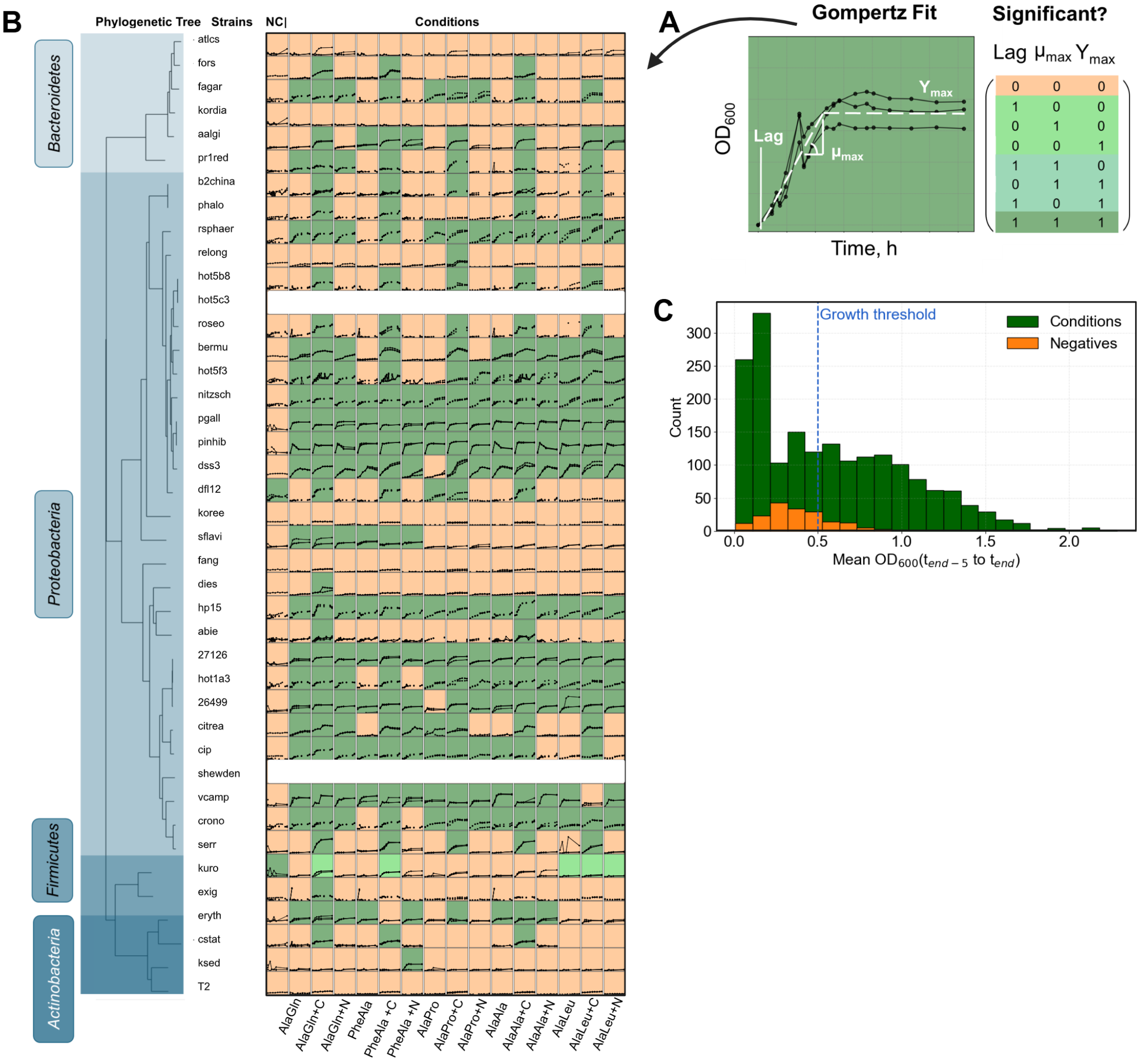


Figure S7: **(A)** Colors indicate the number of parameters that are significant after Gompertz fitting and comparison with growing negative controls. **(B)** Heatmap showing growth on dipeptides, either alone or supplemented with carbon and nitrogen. Note that two strains were not included in this screening. **(C)** Yield data used to define the growth threshold of ln($OD_{600}$) = 0.5.

#### S8 Determining Growth and Growth Parameters

In Figure S8, the workflow used to classify growth versus no growth is illustrated. Measured $OD_{600}$ values were first normalized to the value at time point $t_0$ and subsequently log-transformed. Growth was initially assessed by comparing the mean yield of the final five $OD_{600}$ measurements from each strain and condition to that of the negative controls. Comparing the distribution of the conditions and negative controls resulted in the definition of a growth threshold. Growth data of strains whose yield exceeded the growth threshold (growers) were then fitted with a Gompertz function to estimate growth parameters, including Lag phase ($Lag$), maximum growth rate ($\mu_{max}$), and maximum yield ($Y_{max}$) (see Eq. ES1).

$$OD(t) = Y_{\max} \cdot \exp\left[-\exp\left(\frac{\mu_{\max} \cdot e}{Y_{\max}}\left(\text{Lag} - t\right) + 1\right)\right] \tag{ES1}$$

Strains with a mean yield below the predefined growth threshold were classified as non-gowers and assigned a fixed growth rate of zero, a Lag phase of 50 hours, and the maximum yield observed in the data. For grower strains, the estimated growth parameters were used for subsequent statistical analysis. Specifically, each of the three growth parameters was compared to the corresponding negative control: parameters were considered significant when the estimated growth rate and maximum yield exceeded those of the negative control, or the Lag phase was shorter than the Lag phase of the negative control (see Figure S8).

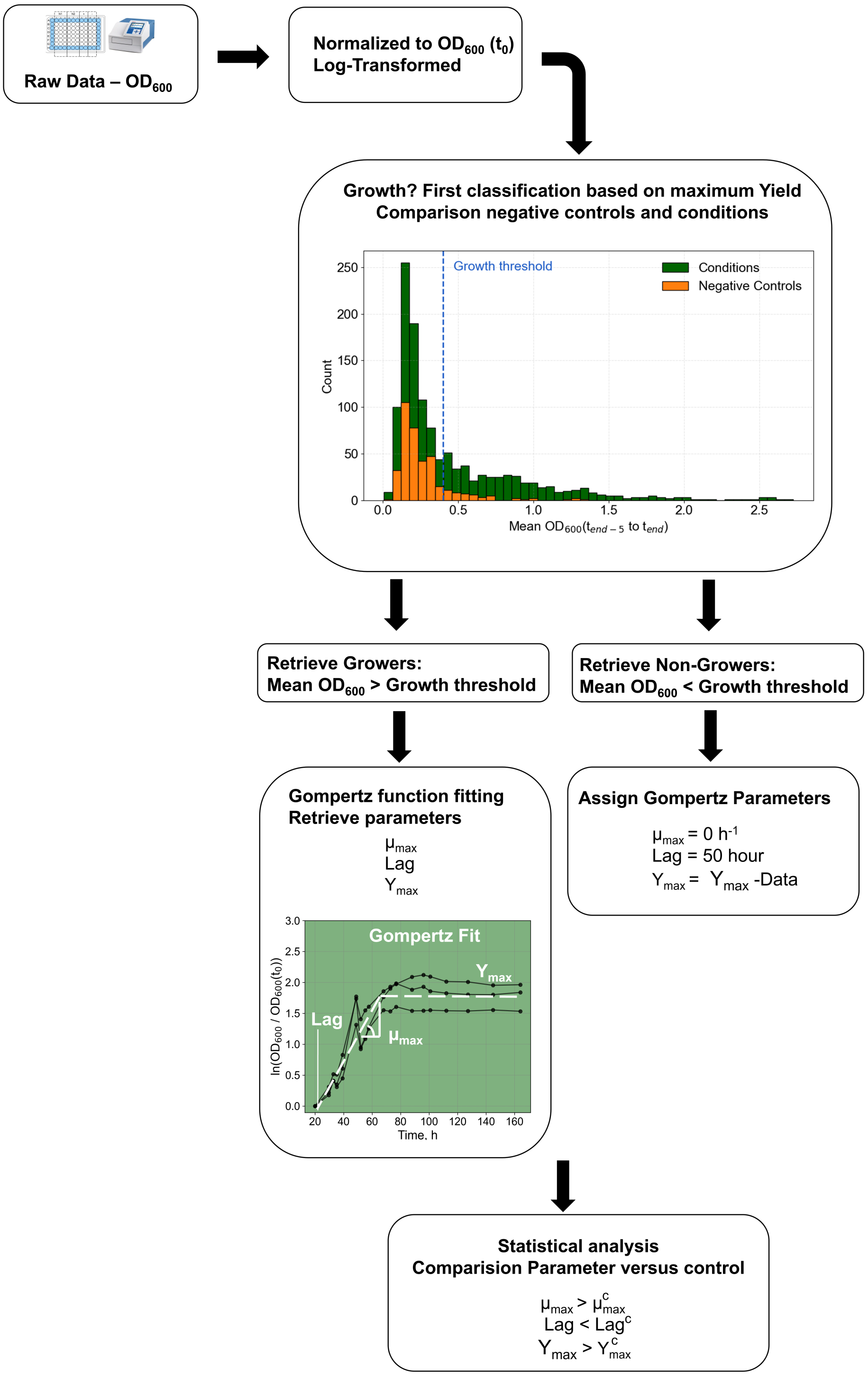


Figure S8: Workflow for classifying growth versus non-growth in screening experiments. Data were normalized and log-transformed. The mean $OD_{600}$ of the final five time points was used to define a growth threshold (blue dashed line) based on comparison with negative controls. Strains with a mean $OD_{600}$ above this threshold were classified as growers and fitted with a Gompertz function to estimate growth parameters, including maximum growth rate (μmax), lag phase, and maximum yield (Ymax). Non-growers were assigned $\mu_{max} = 0$ and a $Lag$ = 50 hours, while maximum yield was taken directly from the data. Growth parameters of growers were subsequently tested statistically against those of growing negative controls (parameter$^{c}$).

#### S9 PCA with additional metadata

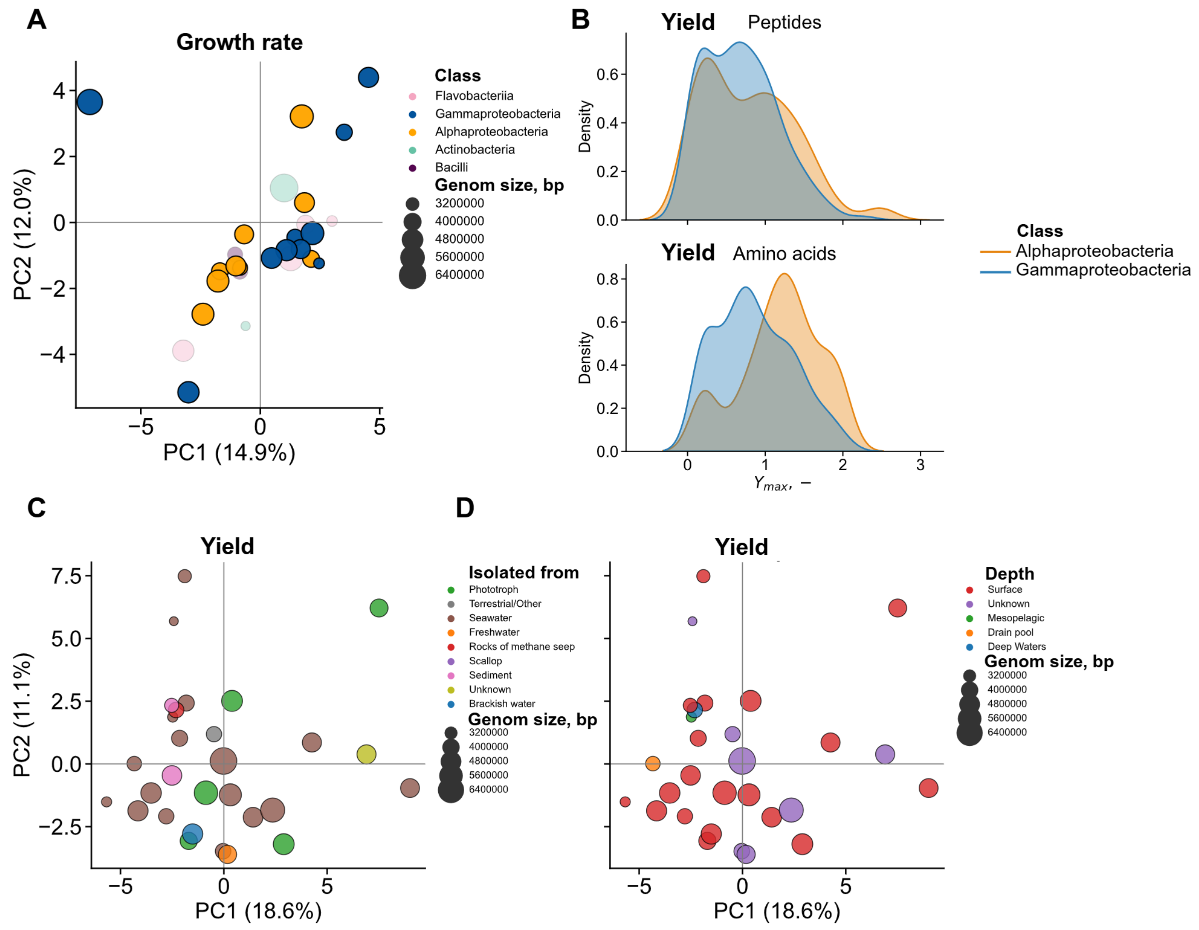


Figure S9: **(A)** Principal component analysis (PCA) of growth-rate phenotypes across experiments E1, E2, and E5. **(B)** Comparison of maximum optical densities (yields) between *Alphaproteobacteria* and *Gammaproteobacteria*. Including additional metadata, such as isolation location **(C)** and sampling depth **(D)**, did not reveal any additional structure within the DIN dataset. See Github for the corresponding supplementary data.

#### S10 Redoxstate as substrate property

Following Chakrawal et al. [5], we calculated the degree of reduction for carbon and nitrogen in neutral molecules as follows:

$$\gamma_C = \frac{4n_C + n_H - 3n_N - 2n_O}{n_C} \tag{1}$$

$$\gamma_N = \frac{5n_N + n_H - 3n_C - 2n_O}{n_N} \tag{2}$$

A combined degree of reduction for carbon and nitrogen was determined as:

$$\gamma_{CN} = \frac{n_c\gamma_c + n_N\gamma_N}{c_C + n_N} \tag{3}$$

Notably, this expression can be rewritten in terms of the molecular C:N ($= \frac{n_C}{n_N}$) ratio:

$$\gamma_{CN} = \frac{C:N\gamma_C + \gamma_N}{C:N+1} \tag{4}$$

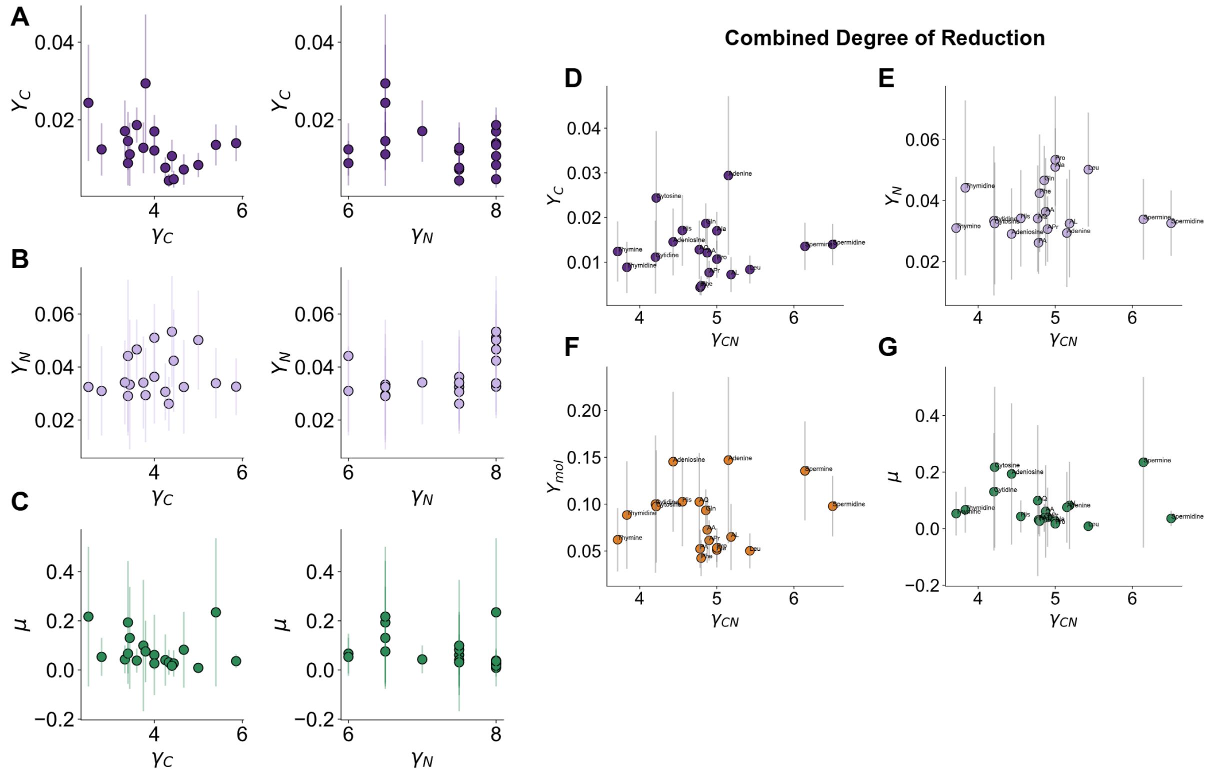


Figure S10: Comparison of normalized yields and growth rates as a function of the degree of reduction of carbon and nitrogen in substrates used as sole carbon and nitrogen sources. Carbon-normalized yield ($Y_C$) **(A)**, nitrogen-normalized yield ($Y_N$) **(B)**, and the non-normalized growth rate **(C)** are shown as a function of the carbon ($\gamma_C$) or nitrogen ($\gamma_N$) degree of reduction. Panels **(D–G)** show carbon-normalized yield ($Y_C$), nitrogen-normalized yield ($Y_N$), substrate-normalized yield ($Y_{mol}$), and growth rate, respectively, as a function of the combined degree of reduction of carbon and nitrogen ($\gamma_{CN}$).

#### S11: Definition of Yields

Given the specific growth rate ($\mu$), the uptake rates of carbon and nitrogen ($r_{C,\mathrm{up}}$ and $r_{N,\mathrm{up}}$), and the rates of carbon and nitrogen incorporation into biomass ($r_{C,\mathrm{bio}}$ and $r_{N,\mathrm{bio}}$), we define the following yields:

$$Y_N = \frac{\mu}{r_{N,\mathrm{up}}} \sim \frac{\mu}{N_{total}}, \tag{5}$$

$$Y_N^{\mathrm{Bio}} = \frac{\mu}{r_{N,\mathrm{Bio}}}, \tag{6}$$

$$Y_C = \frac{\mu}{r_{C,\mathrm{up}}} \sim \frac{\mu}{C_{total}}, \tag{7}$$

$$Y_C^{\mathrm{Bio}} = \frac{\mu}{r_{C,\mathrm{Bio}}}. \tag{8}$$

For the experimental data, uptake rates are not measured directly. Instead, we assume that all supplied substrate is consumed by the end of the experiment, allowing us to approximate uptake using the total amount of carbon and nitrogen supplied, where $N_{total}$ and $C_{total}$ denote the total amounts of nitrogen and carbon provided by each substrate. To provide a conceptual framework for $Y_N$ and $Y_C$, defined as biomass yield per limiting atom, we use *E. coli* as an example. The mass fractions of carbon and nitrogen in biomass are known from Bionumbers (BNID 100649). Using the molar masses of nitrogen and carbon, these can be converted into molar yields:

$$Y_N = \frac{1}{0.14\ gN\ gCDW^{-1}} \cdot \frac{14\ gN\ mmol^{-1}}{1000} = 0.1\ gCDW\ mmol^{-1} \tag{9}$$

The same approach yields the molar carbon yield:

$$Y_C = \frac{1}{0.47\ gC\ gCDW^{-1}} \cdot \frac{12\ gC\ mmol^{-1}}{1000} = 0.026\ gCDW\ mmol^{-1} \tag{10}$$

As a first-order approximation, we assume constant yields and neglect metabolic variability. The quantities $Y_N^{\mathrm{bio}}$ and $Y_C^{\mathrm{bio}}$ denote cell-specific efficiencies and are assumed to remain constant under a fixed biomass C:N ratio. These are expected to exceed $Y_N$ and $Y_C$, respectively.

In flux balance analysis (FBA), growth is represented by the biomass reaction, which is optimized. From the uptake fluxes (exchange reactions, $\nu$) of carbon or nitrogen, biomass production per limiting atom can be calculated as:

$$From\ FBA: Y_N = \frac{\mu}{\nu_N} = [\frac{h^{-1}}{mmol\ gCDW^{-1}\ h^{-1}} = gCDW\ mmol^{-1}] \tag{11}$$

Genome-scale models capture differences in nitrogen and carbon assimilation, although regulatory layers are not explicitly represented. Accordingly, FBA is expected to reflect these differences in the predicted specific growth rate, $\mu$.

**Yields as a function of the C:N ratio**: On the $C : N$ axis we can find a value $\vartheta_{switch}$ at which the limitation switches between carbon and nitrogen. By definition, when $C : N < \vartheta_{switch}$ the cells are C-limited, while at $C : N > \vartheta_{switch}$ the cells are N-limited. Because the total amount of nitrogen supplied was held constant across substrates, increasing substrate C:N ratio corresponds to increasing carbon availability at fixed nitrogen availability. In the carbon-limited regime, additional carbon should therefore increase biomass production, and consequently the nitrogen-normalized yield ($Y_N$) is expected to increase with C:N. Once nitrogen becomes limiting, further increases in carbon availability should no longer increase biomass, and $Y_N$ should approach a plateau. Conversely, the carbon-normalized yield $Y_C$ is expected to remain approximately constant in the carbon-limited regime, where biomass production scales with carbon supply. In the nitrogen-limited regime, biomass is constrained by the fixed nitrogen supply, while carbon supply continues to increase with substrate C:N. As a result, $Y_C$ is expected to decrease with increasing C:N. With this we can define the two functions $Y_C(C : N)$ and $Y_N(C : N)$ accordingly:

$$Y_N(\mathrm{C:N}) = \begin{cases} a\,(\mathrm{C:N}), & \mathrm{C:N} < \vartheta_{\mathrm{switch}} \\ Y_N, & \mathrm{C:N} \geq \vartheta_{\mathrm{switch}} \end{cases} \tag{12}$$

$$Y_C(\mathrm{C:N}) = \begin{cases} Y_C, & \mathrm{C:N} < \vartheta_{\mathrm{switch}} \\ \dfrac{b}{\mathrm{C:N}}, & \mathrm{C:N} \geq \vartheta_{\mathrm{switch}} \end{cases} \tag{13}$$

Using the definitions of the carbon- and nitrogen-normalized yields, the ratio of the limiting yields is equal to the ratio of carbon and nitrogen supplied to the cells (see Equations 5 and 7). Since each substrate is provided as the sole source of both carbon and nitrogen, this ratio corresponds to the substrate C:N ratio:

$$\frac{Y_N}{Y_C} = \frac{r_{C,up}}{r_{N,up}} = \frac{C_{total}}{N_{total}} = C : N_{substrate} \tag{14}$$

Evaluating this expression at the switch point $\vartheta_{switch}$ yields:

$$\frac{Y_N(\vartheta_{switch})}{Y_C(\vartheta_{switch})} = \frac{Y_N}{Y_C} = \vartheta_{switch}. \tag{15}$$

Table 7: Yields per limiting carbon or nitrogen.

| **Derived from** | $\mathbf{Y}_N$ | $\mathbf{Y}_C$ | $\mathbf{Y}_N^{\mathrm{bio}}$ | $\mathbf{Y}_C^{\mathrm{bio}}$ | **C:N Biomass** |
|---|---|---|---|---|---|
| Example | 0.1 | 0.026 | nd | nd | nd |
| FBA simulation *E. coli* | 0.076 $\pm$ 0.023 | 0.014 $\pm$ 0.0021 | 0.093 | 0.024 | 3.8 |
| FBA simulation *A. macleodii* | 0.048 $\pm$ 0.018 | 0.0070 $\pm$ 0.0012 | 0.1 | 0.022 | 4 |
| Data | 0.014 $\pm$ 0.009 | 0.046 $\pm$ 0.018 | nd | nd | nd |

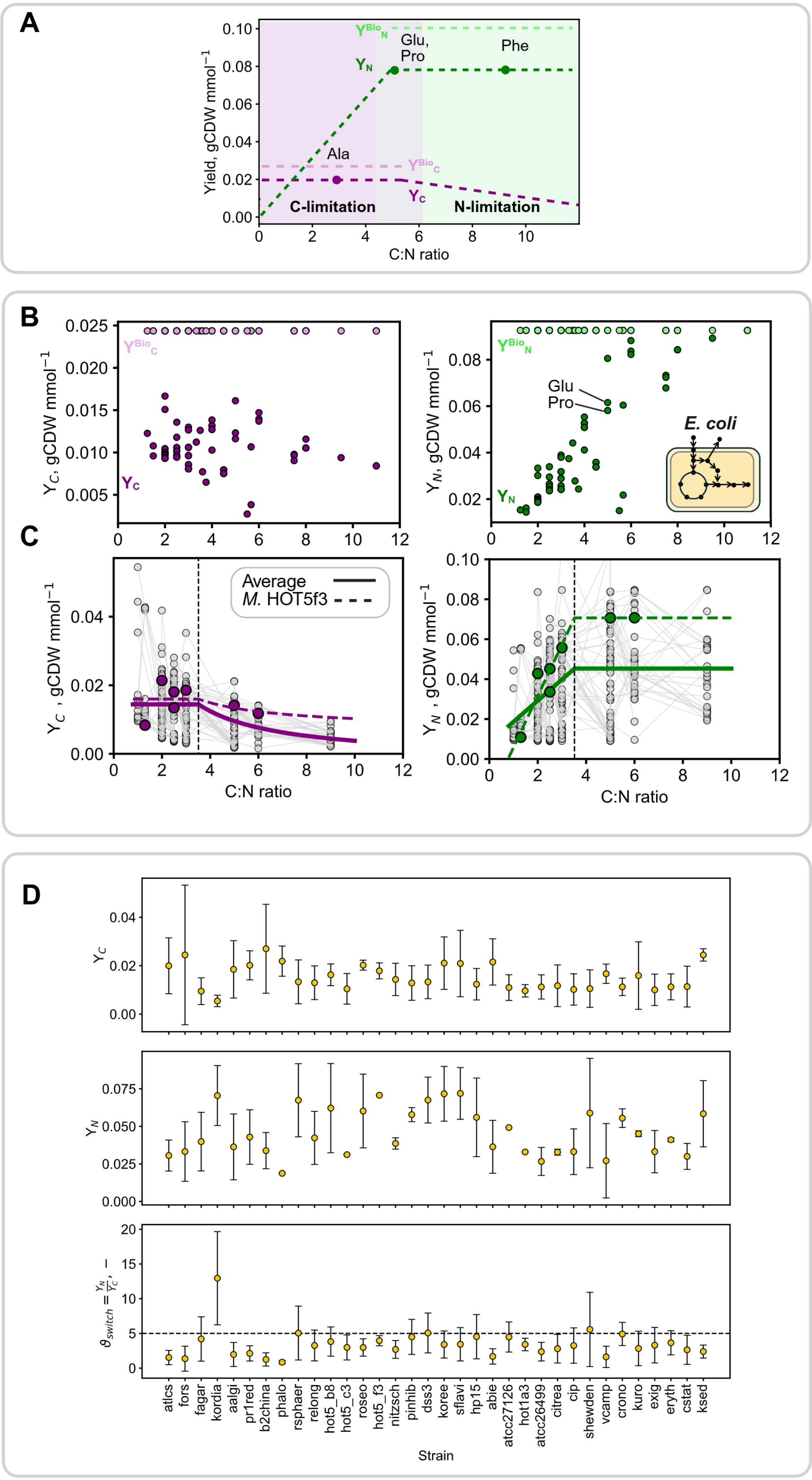
A
Yield, gCDW mmol$^{-1}$
0.10
0.08
0.06
0.04
0.02
0.00
$Y^{Bio}_N$
Glu,
Pro
Phe
$Y_N$
Ala
$Y^{Bio}_C$
$Y_C$
C-limitation
N-limitation
0
2
4
6
8
10
C:N ratio
B
$Y_C$, gCDW mmol$^{-1}$
0.025
0.020
0.015
0.010
0.005
$Y^{Bio}_C$
$Y_C$
$Y_N$, gCDW mmol$^{-1}$
0.08
0.06
0.04
0.02
$Y^{Bio}_N$
Glu
Pro
E. coli
$Y_N$
0
2
4
6
8
10
12
C
$Y_C$ , gCDW mmol$^{-1}$
0.04
0.02
0.00
Average
M. HOT5f3
C:N ratio
$Y_N$ , gCDW mmol$^{-1}$
0.10
0.08
0.06
0.04
0.02
0.00
C:N ratio
D
$Y_C$
0.04
0.02
0.00
$Y_N$
0.075
0.050
0.025
0.000
$\theta_{switch} = \frac{Y_N}{Y_C}$, –
20
15
10
5
0
atlcs
fors
fagar
kordia
aalgi
pr1red
b2china
phalo
rsphaer
relong
hot5_b8
hot5_c3
roseo
hot5_f3
nitzsch
pinhib
dss3
koree
sflavi
hp15
abie
atcc27126
hot1a3
atcc26499
citrea
cip
shewden
vcamp
crono
kuro
exig
eryth
cstat
ksed
Strain

#### S12 Relationship between $Y_c$ and $Y_N$

Figure S12: **(A)** Relationship between $Y_N$, $Y_C$, and the substrate C:N ratio. $Y_C$ and $Y_N$ are constant under carbon- and nitrogen-limited conditions, respectively. $Y_C$ decreases at higher C:N ratios due to carbon-replete conditions, whereas $Y_N$ decreases at lower C:N ratios, where carbon availability limits biomass production (note that the decrease in $Y_C$ is hyperbolic). The quantities $Y_C^{\mathrm{bio}}$ and $Y_N^{\mathrm{bio}}$ represent the growth relative to the rate of carbon or nitrogen incorporated into biomass (see Equations (2) and (4)). **(B)** FBA simulation of the *E. coli* model iML1515 showing $Y_N$ and $Y_C$ across a range of C:N ratios. $Y_C^{\mathrm{bio}}$ and $Y_N^{\mathrm{bio}}$ remain constant and can be obtained from the biomass reaction of the model or calculated using Equations (2) and (4). **(C)** Experimentally obtained mean $Y_N$ and $Y_C$ as a function of the C:N ratio (solid lines), with an example shown for *Marinovum* HOT5f3 (dashed lines). **(D)** Yields across strains and $\vartheta_{\text{switch}}$, estimated from the ratio of $Y_N$ and $Y_C$.

#### S13: Definitions of nitrogen use efficiency (NUE)

Carbon and nitrogen incorporation rates are defined as follows, with $r_{N,\mathrm{Sec}}$ and $r_{C,\mathrm{Sec}}$ representing the respective secretion rates.

$$r_{N,\mathrm{Bio}} = r_{N,\mathrm{up}} - r_{N,\mathrm{Sec}}, \tag{16}$$

$$r_{C,\mathrm{Bio}} = r_{C,\mathrm{up}} - r_{C,\mathrm{Sec}}. \tag{17}$$

Following the definition of [6] we can express nitrogen use efficiency as:

$$\mathrm{NUE} = \frac{r_{N,\mathrm{Bio}}}{r_{N,\mathrm{up}}} = \frac{r_{N,\mathrm{up}} - r_{N,\mathrm{Sec}}}{r_{N,\mathrm{up}}} = 1 - \frac{r_{N,\mathrm{Sec}}}{r_{N,\mathrm{up}}}. \tag{18}$$

Because neither $r_{N,\mathrm{Bio}}$ nor $r_{N,\mathrm{up}}$ can be easily measured for a variety of DON substrates, we can estimate NUE indirectly. Specifically, we use the biomass yield per nitrogen ($Y_N$) as a function of the substrate C:N ratio, $\vartheta$, derived from the DON screening (E2). Together with the specific growth rate ($\mu$) and the nitrogen secretion rate ($r_{N,\mathrm{Sec}}$), this allows us to approximate the rate of nitrogen assimilation into biomass (see Supplement S12 for details):

$$r_{N,\mathrm{up}} = \frac{\mu}{Y_N(\vartheta)}. \tag{19}$$

$$\mathrm{NUE}(\vartheta) = 1 - \frac{r_{N,\mathrm{Sec}} \cdot Y_N(\vartheta)}{\mu}. \tag{20}$$

Another alternative formulation is:

$$\mathrm{NUE}_{Bio} = \frac{\mu}{\mu + Y_N^{\mathrm{Bio}} \cdot r_{N,Sec}}. \tag{21}$$

In FBA, all fluxes are accessible, enabling calculation of different formulations of NUE, which yield equivalent results (see Figure S13). Furthermore, as described in [6], NUE and CUE are coupled through substrate stoichiometry and the biomass C:N ratio:

$$r_{N,\mathrm{Bio}} = \mathrm{NUE} \cdot r_{N,\mathrm{up}}, \tag{22}$$

$$r_{C,\mathrm{Bio}} = \mathrm{CUE} \cdot r_{C,\mathrm{up}}, \tag{23}$$

$$\frac{C}{N}_{biomass} = \frac{r_{C,\mathrm{Bio}}}{r_{N,\mathrm{Bio}}} = \frac{\mathrm{CUE} \cdot r_{C,\mathrm{up}}}{\mathrm{NUE} \cdot r_{N,\mathrm{up}}} = \frac{\mathrm{CUE}}{\mathrm{NUE}} \cdot \frac{C}{N}_{substrate}. \tag{24}$$

We

Table 8: Calculation of NUE

| **Condition** | **Data** | **Equation used** |
|---|---|---|
| *E. coli* FBA siumlation (iMl1515) | Simulated growth rate on DON | [20] |
| *A. macleodii* FBA siumlation (iHS4156) | Simulated growth rate on DON | [20] |
| Marine heterotrophic strains | Yield obtained from DON | [21] |

### S14 Nitrogen use efficiency for *Alteromonas macleodii* MIT

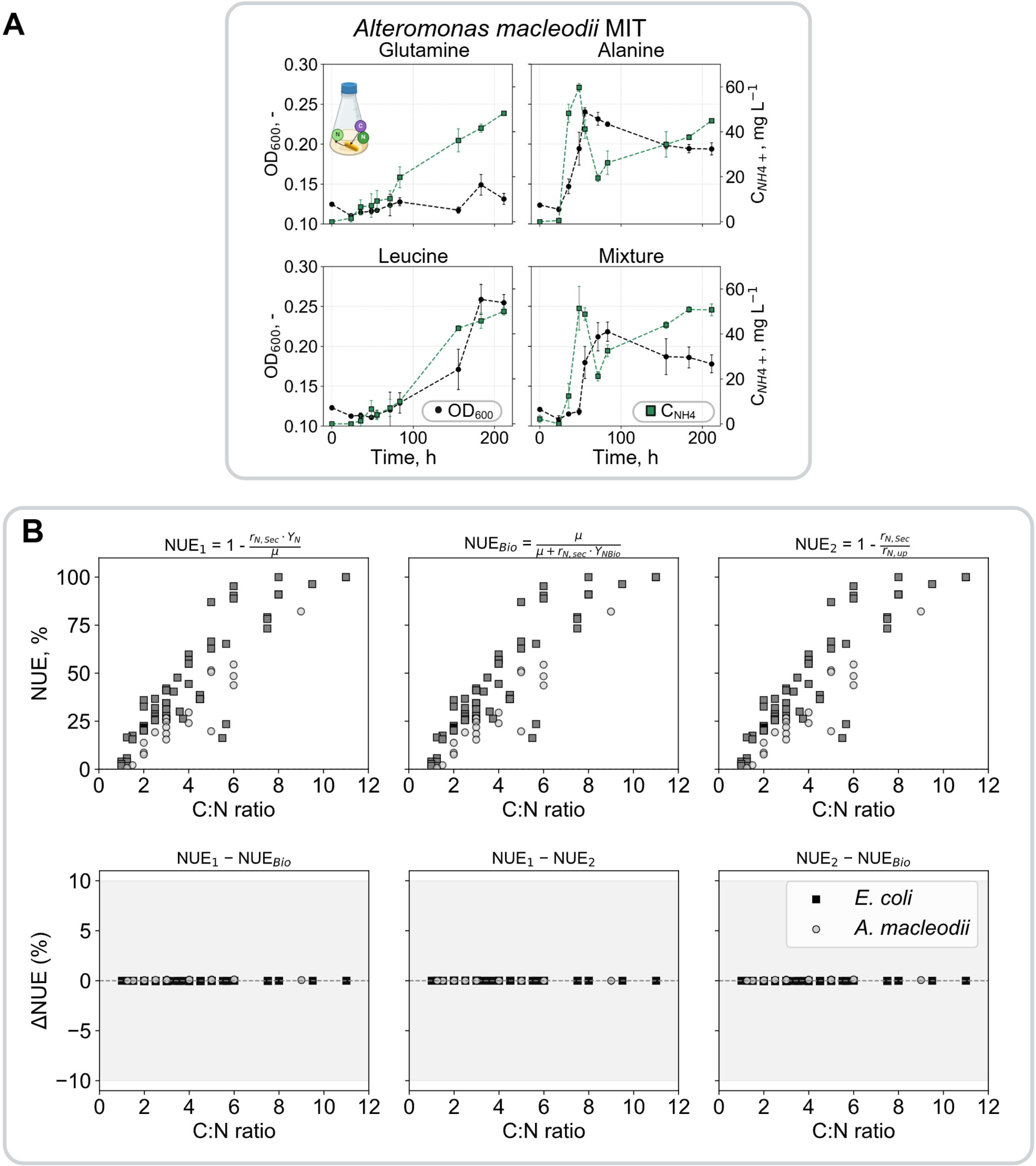


Figure S14: **(A)** Example of growth experiment for determining NUE. **(B)** Comparison of different NUE expressions calculated from FBA simulations of the *E. coli* model. All expressions are equivalent when using fluxes obtained from the simulations.

#### S15 Non-linear effects for inorganic compounds

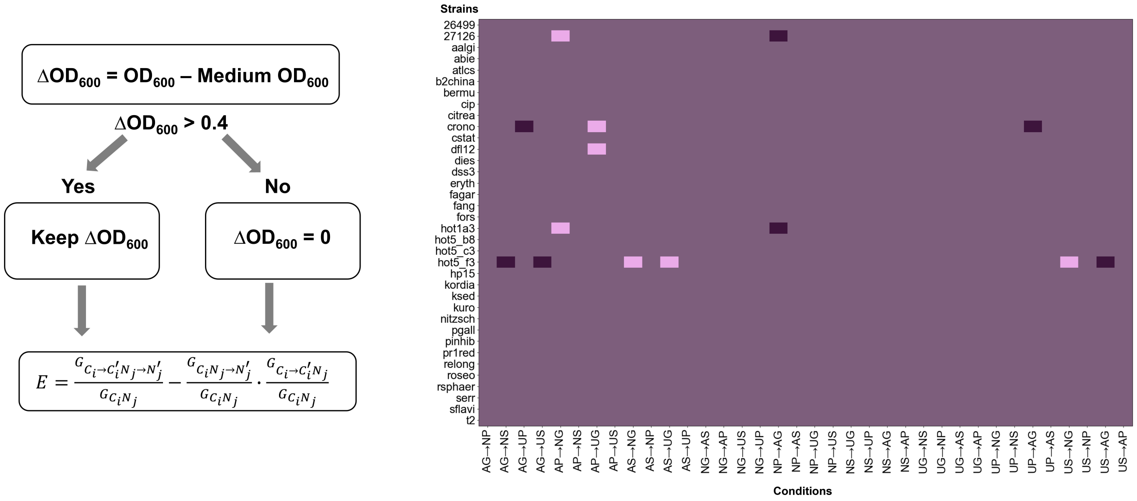


Figure S15: Calculation of epistasis. $OD_{600}$ values above the growth threshold (0.4) were used for epistasis calculations. The heatmap shows outliers, defined as values exceeding ±1 standard deviation of the epistasis distribution across all strains and conditions (light purple: above; dark purple: below).

#### S16: Epistasis may be dependent on nutrient limitation

Because our DIN screening focused on nitrogen effects, epistasis was analyzed under nitrogen-limited conditions (C:N = 20). Using the *E. coli* genome-scale model iML1515, we simulated growth on the same carbon and nitrogen sources across a range of C:N ratios to capture both carbon limitation (low C:N) and nitrogen limitation (high C:N) (see Supplementary Figure S8). We found that growth responses are nearly identical across nutrient combinations (nitrate or ammonium paired with glucose, succinate, or pyruvate) under nitrogen limitation, but become more distinct under carbon limitation. This is consistent with the minimal epistasis observed in experimental yields under nitrogen limitation and suggests that the observed outliers may reflect strain-specific differences in nitrogen metabolism.

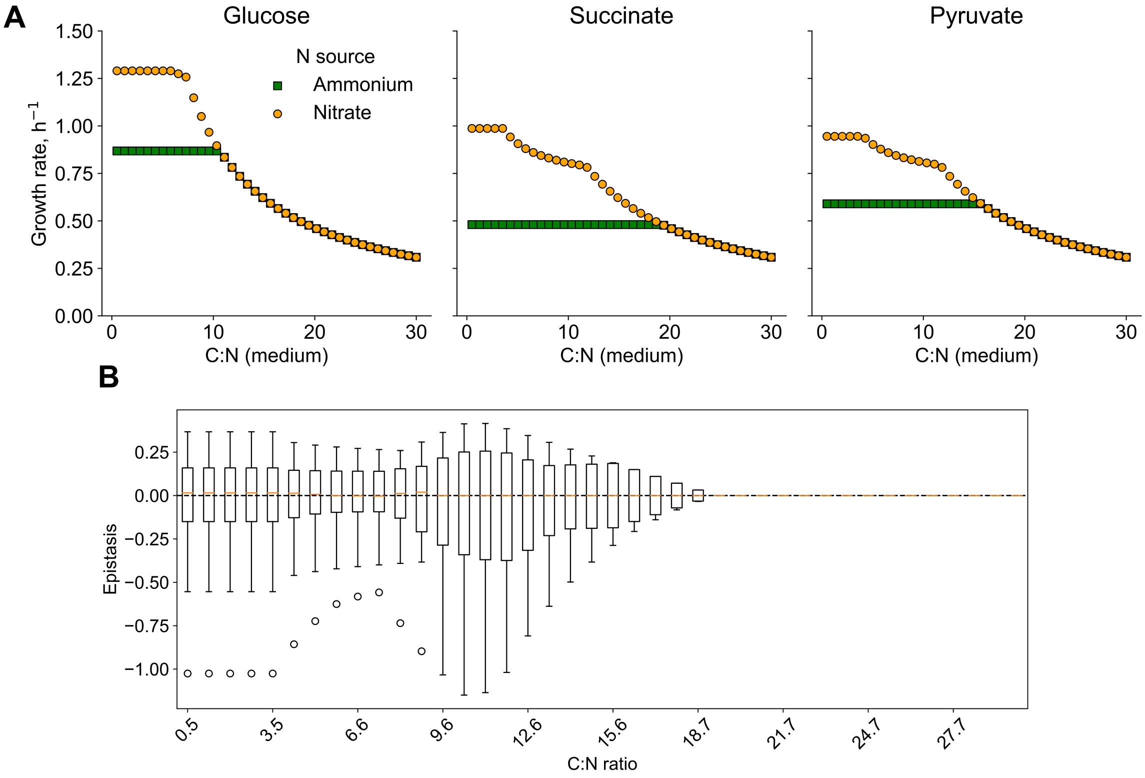


Figure S16: **(A)** Growth simulations of *E. coli* iML1515 on glucose, succinate, and pyruvate as sole carbon sources, with either ammonium (green squares) or nitrate (orange circles), across a range of C:N ratios. Carbon uptake was fixed at 100 mmol gCDW$^{-1}$ h$^{-1}$, while nitrogen availability was varied to achieve different C:N ratios. **(B)** Epistasis distributions across C:N ratios. Distributions are centered near zero at higher C:N ratios, indicating minimal epistasis under nitrogen limitation.

#### S17: Details on normalization:

Experimentally, amino acid mixtures were normalized to a total nitrogen concentration of 10 mM. In each mixture, all amino acids contributed equally to the total nitrogen (e.g., in a mixture of three amino acids, each contributed 3.33 mM N). To account for differences in carbon supply, yields were additionally normalized by the amount of carbon provided (i.e., divided by moles of carbon). Because we expected cells to be nitrogen-limited on proline and leucine, we used the maximum usable carbon concentration for Leucine and a mixture of leucine and proline (50 mM*) for normalization. Table 9 shows the provided nitrogen and carbon and the used normalization.

For each amino acid mixture, the observed yield was compared to the mean yield of the corresponding individual amino acids measured for the same strain.
The fold change was calculated as:

$$\mathrm{FC}_{\mathrm{raw}} = \frac{\text{Observed OD}_{600}}{\text{Expected OD}_{600}}$$

where

$$\text{Expected OD}_{600} = \frac{1}{n}\sum_{i=1}^{n} \text{OD}_{600,i}$$

is the mean $OD_{600}$ obtained from the individual amino acids present in the mixture.

Table 9: Carbon and nitrogen supplied for amino acid combinations. (*marks normalization cut at 50 mM); A = adenine, Q = glutamine, P = proline, L = leucine.

| Condition | N per AA (mM) | Total C (mM) |
|---|---|---|
| *Single amino acids* | | |
| A | 10 | 30 |
| Q | 10 | 25 |
| P | 10 | 50 |
| L | 10 | 60* |
| *Pair combinations* | | |
| AQ | 5 | 27.5 |
| AL | 5 | 50 |
| AP | 5 | 40 |
| QL | 5 | 47.5 |
| QP | 5 | 37.5 |
| LP | 5 | 55* |
| *Triple combinations* | | |
| AQL | 3.33 | 38.33 |
| AQP | 3.33 | 35 |
| ALP | 3.33 | 46.67 |
| QLP | 3.33 | 45 |
| *Quadruple combination* | | |
| AQLP | 2.5 | 41.25 |

#### S18 Comparison between carbon normalized and raw fold changes

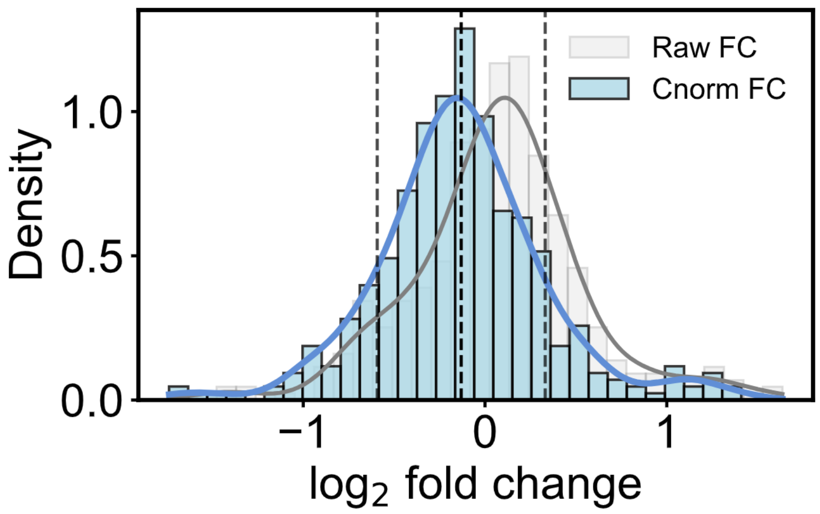


Figure S18: Comparison of fold changes (FC) calculated from carbon-normalized and raw data for growth on different amino acid mixtures.
